# Complete genome-derived metabolic interactions reveal impact of gut ecology on human health

**DOI:** 10.64898/2026.03.25.714329

**Authors:** Yuzheng Gu, Haoyu Wang, Jinlong Yang, Tao Zeng, Hewei Liang, Wenxin He, Mengmeng Wang, Zhinan Wu, Liye Yang, Yu-sheng Xu, Juan Zhao, Yuning Zhang, Yuliang Dong, Yiyi Zhong, Haifeng Zhang, Jinhong Wang, Xing Rao, Yangfeng Wen, Xiaofan Sun, Karsten Kristiansen, Yu Tian, Xin Tong, Ya Wang, Juan Yang, Fushu Liu, Zejun Yang, Wangsheng Li, Bo Wang, Peng Gao, Jun Xu, Yinglei Miao, Xin Jin, Chuanyu Liu, Xun Xu, Yang Sun, Feng Zhao, Liang Xiao, Yuanqiang Zou

## Abstract

Metabolic interactions govern gut microbiome assembly, yet functional rules remain obscured by genomic incompleteness and fragmentation. Here, we leverage 1,150 complete genomes to construct genome-scale metabolic models, demonstrating that draft genomes introduce systemic artifacts missing critical transport functions. We observe that metabolic interactions are intrinsically driven by genomic traits and niche specialization rather than random association. We stratifies strains into four ecological groups, including active players, resource predators, utilizers, and contributors, each of which exhibits distinct metabolite exchange/competition signatures. This functional architecture is strongly coupled to secondary metabolism. Applying this framework to inflammatory bowel disease revealed distinct temporal dynamics of ecological groups across disease subtypes, where group-specific dysbiosis predicted clinical phenotypes better than whole community profiles. Furthermore, integrating metabolic and co-occurrence networks to identify keystone for classifiers as features significantly improved disease diagnosis. Our study provides novel insights into gut metabolic interaction landscape, bridging the gap between genomic potential and clinical utility for precision microbiome therapeutics.

## Introduction

The gut microbiota functions as a pivotal metabolic organ, regulating host physiological homeostasis in part through the synthesis of bioactive compounds. Microbial metabolites, such as short-chain fatty acids, bile acids, and vitamins, serve as essential mediators that modulate host immunity, energy metabolism, and neural signaling^1-5^. While the physiological effects of these metabolites have been extensively characterized, the ecological mechanisms governing their production, specifically, how microbial community members coordinate their metabolic activities remains poorly understood^4,6,7^. A systematic exploration of this coordination is essential to elucidate the rules of community assembly and their ultimate impact on host health.

Metabolic activities within the gut are not isolated events but are driven by complex interactions, including nutritional competition, cross-feeding, and chemical communication^8-10^. Although existing research indicates that anomalous interaction patterns are closely associated with diseases such as inflammatory bowel disease (IBD)^11^, obesity^12^, and colorectal cancer^13^, current inferences about microbial interactions mainly rely on abundance-based association analyses^14,15^. Metabolic dependencies (e.g. cross-feeding) between microbes and substrate competition are the essential driving forces shaping microbial community structure^16,17^. For instance, Bacteroides species can degrade various complex polysaccharides to produce oligosaccharides, providing growth substrates for Bifidobacterium species^18^, while competition for iron ions between *Escherichia coli* and *Salmonella enterica* influences the colonization of intestinal pathogens^19^. However, statistical co-occurrence often reflects shared environmental preferences rather than genuine metabolic dependencies^20,21^. Consequently, identifying key functional taxa and translating them into diagnostic biomarkers based on the underlying logic of metabolic interactions remains a significant challenge.

Genome-scale metabolic models (GEMs) offer a mechanistic alternative by simulating metabolic fluxes to predict cross-feeding and competition^22,23^. However, a critical bottleneck restricts the current field, namely that the vast majority of GEMs are reconstructed from draft genomes or metagenome-assembled genomes (MAGs)^24,25^. Metabolic interactions rely strictly on the exchange of molecules, which is mediated by transporters and secretion systems^26,27^. The genes involved in these interactions are often located in genomic regions that are prone to fragmentation in draft assemblies. Consequently, GEMs derived from incomplete genomes frequently miss critical transport reactions, leading to severe biases or false negatives in predicting metabolic exchanges^28,29^. To accurately reconstruct the interactive network of the microbiome, GEMs based on high-quality genomes are crucial^30^.

Here, we bridged this gap by constructing 1,150 high-quality GEMs using complete genomes isolated from healthy individuals. Leveraging this dataset, we demonstrate that complete genomes recover a vast number of transport reactions overlooked by draft-based models, thereby enabling a high-resolution quantification of metabolic competition and complementarity. Based on the asymmetry of these interactions, we established a novel ecological framework that stratifies gut bacteria into four distinct groups: active players, resource predators, resource utilizers, and resource contributors. Furthermore, by integrating these metabolic interaction features with clinical data, we reveal that keystone taxa identified through metabolic networks significantly enhance the performance of disease diagnostic models. This study provides a genome-resolved ecological framework for understanding gut microbiota assembly (Supplementary Fig. S1) and offers new avenues for precision diagnostics based on metabolic mechanics.

## Results

### Advantages of constructing GEMs using complete genomes

To assess the improvement of complete genomes for constructing GEMs, we selected 966 models developed from complete genomes (Complete GEMs) and their corresponding counterparts constructed from draft genomes based on the same strain for comparative analysis. Although our analysis focused on 199 species, these taxa demonstrated extensive prevalence and consistent abundance across diverse cross sectional cohorts (China, HMP, and the Netherlands), indicating a high coverage of the core gut microbiota (Supplementary Fig. 2). We first quantified the genomic differences and found that complete genomes as expected contained a significantly higher number of unique coding sequences (CDS) (49,288) compared to draft genomes (6,862) (Fig. 1A). Functional characterization revealed that genes unique to complete genomes were enriched in mobile elements like transposases, whereas CDS unique to draft genomes were significantly shorter in length (Fig. 1B) and enriched in low-quality fragments such as uncharacterized proteins. This length discrepancy suggests that many draft-unique genes are likely assembly artifacts or truncated fragments rather than genuine contiguous biological sequences. This genomic fragmentation directly affected the accuracy of metabolic reconstruction. While the majority of metabolic reactions (average 1,507) were shared between paired models (Fig. 1C), we observed a significant positive correlation (R = 0.18, P = 0.006) between the number of draft-unique CDS and the number of unique reactions in draft-based models (Fig. 1D). This indicates that the fragmented CDS in draft assemblies often led to the inclusion of spurious reactions (false positives) in the models. In contrast, reactions unique to complete genome-based GEMs represented genuine metabolic capabilities that were overlooked in draft reconstructions. We also compared our generated genome-based metabolic models with AGORA2 and Apollo^31,32^. Here we observed that, in terms of reactions, the average number of reactions of the Complete GEMs (1506.69 ± 530.51) was close to that of the manually processed AGORA2 (1723.12 ± 817.14), and far exceeded that of the automatically processed Apollo (997.92 ± 215.34) (Supplementary Fig. S3).

**Fig. 1.**
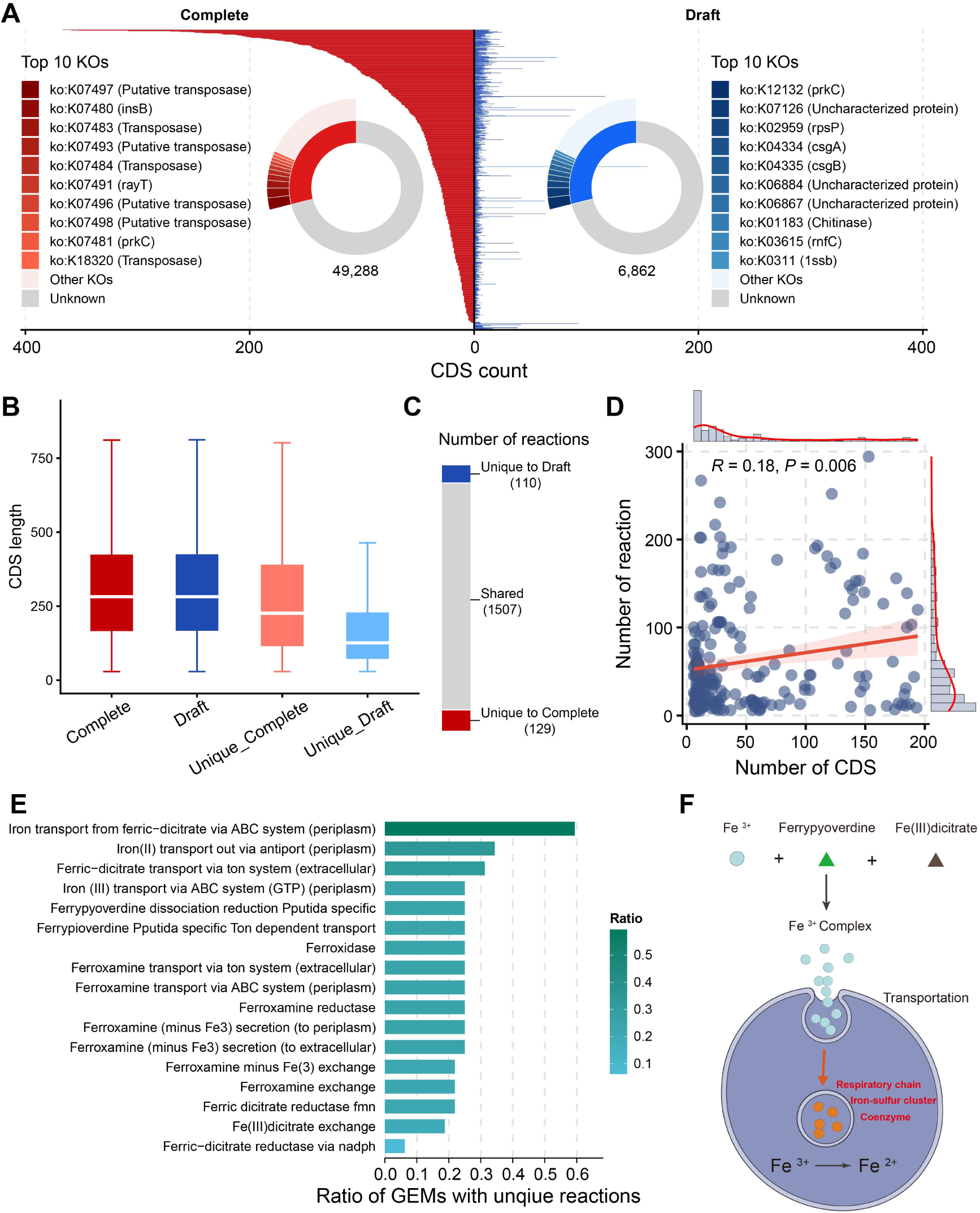
The advantages of genome-scale metabolic models (GEMs) constructed from complete genomes compared with draft genomes. **A** Quantitative and functional discrepancy of coding sequences (CDS) between complete and draft genome assemblies. Barplots displays the count of CDS unique to complete and draft genomes for each strain. Donut charts illustrate the characterization of these unique genes: the inner ring represents the proportion of CDS with functional annotations, while the outer ring highlights the top 10 most prevalent KEGG Orthologs (KOs). **B** Comparison of CDS length distributions across four categories, including total CDS in complete and draft genomes, and CDS unique to each assembly type. **C** Composition of GEMs reconstructed from paired assemblies. The stacked bar shows the average proportion of metabolic reactions shared between assemblies, and those unique to complete and draft genome-based models. **D** Correlation analysis showing the positive association between the number of draft-unique CDS and the number of potential spurious reactions (unique reactions in draft genome-based models) added to the models. **E** Recovery of overlooked metabolic reactions in complete genome-based models. The bar chart displays the prevalence (percentage of models) of specific iron transport and metabolism reactions identified in complete *Lactococcus* models but missed in draft-based reconstructions. **F** A diagram showing how ferric ions (Fe^3+^) bind to cofactors, enter bacteria, and are converted into ferrous ions (Fe^2+^) to participate in biological reactions.

Paired comparisons of GEMs reconstructed from draft and complete genomes for each strain revealed a notable variation in Jaccard dissimilarity of metabolic reactions (median = 0.13) (Fig. S4A). We identified 4,764 additional reactions in the GEMs from the complete genomes compared to draft genomes (Supplementary Table 4), with a majority of these being related to transport. Additionally, other reactions such as those catalyzed by glutamate decarboxylase and prephenate transaminase, were prominently observed within the genera *Escherichia* and *Collinsella* in GEMs from complete genomes but absent inferred from draft genomes. Metabolic reactions of *Lactococcus* based on the complete genome were more numerous than those observed based on the draft genome (Fig. S4B). Specifically, models built by complete genomes of *Lactococcus* significantly improved the recovery of transport systems, particularly processes involved in iron metabolism (Fig. 1E). Key reactions involved in the uptake of ferric-dicitrate and ferroverdin were prevalent in complete models but largely absent in drafts. These recovered pathways are critical for capturing the full mechanism of iron acquisition, where ferric ions (Fe^3+^) are bound by cofactors, transported into the cell, and reduced to ferrous ions (Fe^2+^) to participate in essential biological processes (Fig. 1F). Collectively, these results demonstrate that using complete genomes not only reduces the noise from assembly artifacts but also restores critical metabolic functions related to niche adaptation.

### Effects of genomic and phylogenetic architectures on the landscape of metabolic interactions

Insights into microbial metabolic interactions offer a mechanistic pathway to modulate microbiome composition and host physiology^33^. To quantify these interactions, we calculated metabolic competition and complementarity indices between all strain pairs. The competition index reflects the potential overlap in nutritional requirements, whereas the complementarity index quantifies the capacity of one organism to synthesize metabolites required by another. Due to the asymmetries of indices (Supplementary Fig. 5), a genome could be both the metabolic giver and taker for complementarity and be both the competitor and victim for competition^34^. For instance, *Megasphaera sp000417505* was found to take up fumarate secreted by *Bifdobacterium adolescent* (Supplementary Fig. 6), while also released ammonia as compensation, which has been confirmed in a previous study^35^. Given the significant negative correlation observed between metabolic competition and complementarity indices (Fig. 2A), we integrated them into the metabolic distance score, calculated as 1 - (competition index - complementarity index), where a smaller distance represented higher niche overlap or preference^36^ (Fig. 2B).

**Fig. 2.**
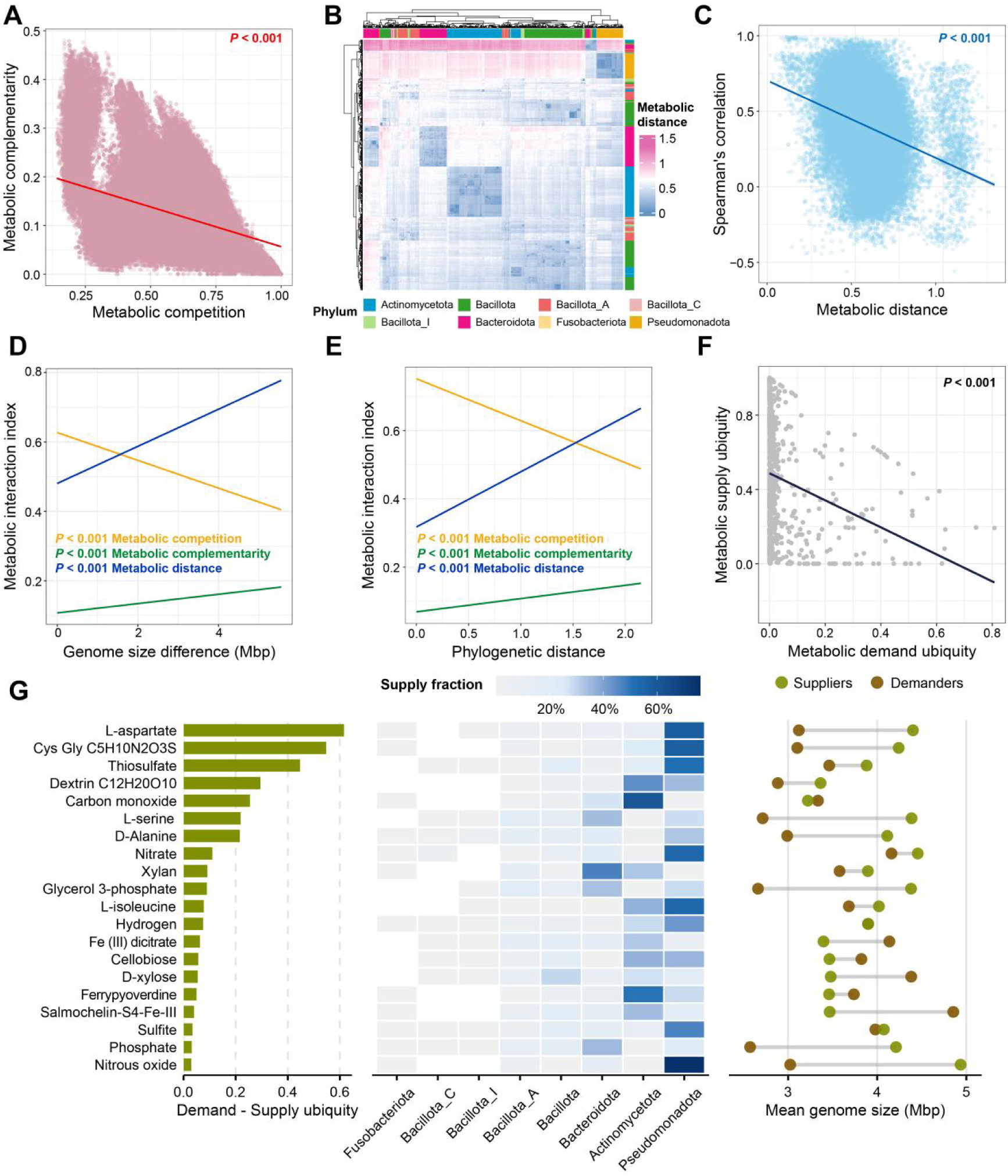
Genomic and phylogenetic constraints govern the pattern of metabolic interactions. **A** Relationship between metabolic complementarity and competition indices derived from GEMs. **B** Heatmap visualization of metabolic distances between pairs of strains, quantifying the metabolic preferences between distinct bacteria. **C** Validation of model-predicted interactions against real-world abundance data. Genomes were mapped to a healthy human cohort metagenomic dataset to calculate pairwise Spearman’s correlations of species abundance. The scatter plot correlates these empirical co-occurrence patterns with their calculated metabolic distances, assessing whether metabolic similarity drives ecological exclusion or co-existence. Relationship of metabolic competition, complementarity, and distance with genome size difference (**D**) and phlylogenetic distance (**E**). **F** Correlation analysis between the supply ubiquity (frequency of producers) and demand ubiquity (frequency of consumers) for all metabolites. **G** The barplot on the left ranks the top 20 metabolites with the highest scarcity scores (defined as demand ubiquity - supply ubiquity). Heatmap on the middle shows the phylogenetic distribution of suppliers for these scarce resources. Dumbbell plot displays the genomic potential gradient comparing the mean genome size of suppliers versus consumers.

To investigate whether metabolic similarity drives ecological exclusion or co-existence, we validated the metabolic distance against empirical abundance data from human cohorts of healthy individuals. We observed a significant negative correlation between metabolic distance and species co-occurrence (Fig. 2C), consistent with previous findings on *Enterobacteriaceae*^36^. This suggests that habitat filtering, where environmental factors select for organisms with shared metabolic traits, overrides the effects of resource competition in shaping the gut microbiome assembly. We next investigated the evolutionary and genomic drivers underlying these interaction patterns. Metabolic distance was strongly positively associated with phylogenetic distance, confirming that metabolic capabilities were phylogenetically conserved (Fig. 2D). Closely related strains share conserved metabolic networks, leading to high niche overlap, whereas distantly related strains possess divergent biosynthetic repertoires that foster complementarity. Furthermore, metabolic interactions were significantly shaped by genome size disparities (Fig. 2E). Specifically, strain pairs with larger differences in genome size exhibited higher complementarity. This observation supported the "Black Queen Hypothesis" and reductive evolution as key drivers of cross-feeding, where organisms undergoing genome streamlining become metabolically dependent on versatile and large-genome community members^37^. Additionally, linear mixed-effect models controlling for phylogeny and genome size revealed that differences in cell wall/membrane/envelope biogenesis genes were the strongest predictor of metabolic interaction indices. Larger disparities in these gene proportions were associated with reduced competition and enhanced complementarity (Supplementary Fig. 7 and Supplementary Table 8).

Finally, we observed a striking negative correlation between metabolic supply ubiquity (proportion of producers) and demand ubiquity (proportion of consumers) (Fig. 2F). This inverse relationship indicated that resources required by the majority of the community were produced by only a limited subset of strains, reflecting widespread metabolic specialization. Focusing on the top 20 most scarce metabolites defined by the highest disparity between demand and supply, we found that their production was not randomly distributed but monopolized by specific taxa, primarily within the phyla Pseudomonadota and Actinomycetota (Fig. 2G). Crucially, for most scarce metabolites (e.g., L-aspartate), the genomes of suppliers were obviously larger than those of demanders (Fig. 2G). Conversely, demanders of specific metabolic byproducts possessed larger genomes, which indicates that they have adopted a strategy of exploiting public goods or waste products released by streamlined strains.

### Distinct ecological groups shaped by metabolic interactions

To systematically decode the functional architecture of the gut microbiome, we stratified 1,150 strains into four distinct ecological groups based on the directionality and asymmetry of their metabolic interactions, including active players, resource predators, resource utilizers, and resource contributors (Fig. 3A, See details in ***Methods***). The robustness of this classification was rigorously validated via bootstrap resampling (n = 100 iterations), yielding an average consistency score > 0.9 for each group (Supplementary Fig. 8A). While we observed a broad association between these ecological groups and taxonomic lineages, the two dimensions were not identical. For instance, while the phylum Pseudomonadota was predominantly mapped to resource contributors, other phyla were distributed across all four groups (Fig. 3B). Also, different strains of the same species (e.g., *Phocaeicola vulgatus* and *Bifidobacterium pseudocatenulatum*) could be assigned to distinct ecological groups (Fig. 3B), suggesting the high diversity and heterogeneity at the strain level.

**Fig. 3.**
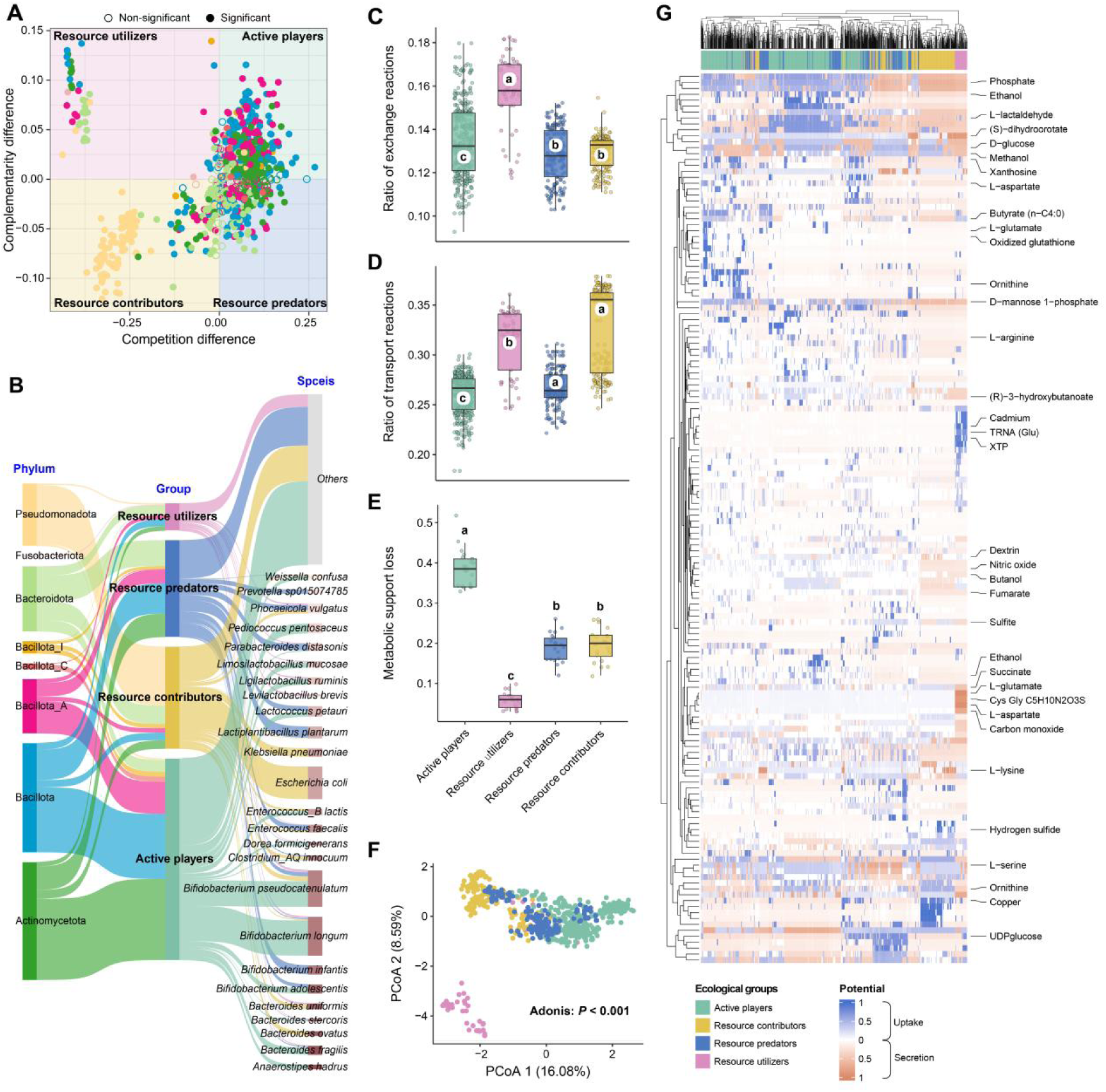
The ecological shaping effect of metabolic interactions on gut bacteria. **A** The division of strains into four ecological taxa including active players, resource predators, resource utilizers, and resource contributors, based on their metabolic interaction directionality. **B** Taxonomic composition of the identified ecological groups. Stacked bars display the distribution of strains at the phylum and species levels within each group. **C** Boxplots showing the proportion of exchange reactions (**C**) and transport reactions (**D**) across the four groups. **E** Impact of targeted groups removal on community metabolic stability. Lowercase letters represent the significance of differences among four groups. Random communities (n = 20) were simulated with 100 strains each. **F** Principal Coordinate Analysis (PCoA) of metabolic complementarity potential profiles for ecological groups. **G** Heatmap illustrating the specific metabolic complementarity potential of different ecological groups. The color scale indicates the potential, defined as the proportion of the total community with which a focal strain uptakes/secretes for a specific metabolite.

We next characterized the metabolic and genomic characteristics underlying these groups (Fig. 3C and D, Supplementary Fig. 8B). Active players exhibited the lowest proportion of transport and exchange reactions concomitant with the smallest genome sizes, suggesting a physiology governed by metabolic streamlining and efficient intracellular processing. In contrast, resource utilizers maintained high proportions of transport and exchange reactions, reflecting a heavy reliance on absorbing environmental resources. Resource contributors possessed the largest genomes and displayed the highest transport capacity, reflecting a robust physiological potential for active metabolite secretion. To validate the ecological significance of these groups, we performed *in silico* perturbation experiments on randomized synthetic communities (Fig. 3E, see details in ***Methods***). Active players emerged as the most critical structural members and their removal precipitated a collapse in community metabolic support (i.e., stability loss), significantly exceeding the impacts of other groups. Conversely, communities displayed high resilience to the loss of resource utilizers, confirming their role as peripheral beneficiaries that rely on cross-feeding fluxes without being structurally critical for community survival.

Principal Coordinate Analysis (PCoA) of metabolite exchange profiles revealed distinct signatures (Fig. 3F). For instance, succinate and carbon monoxide could be secreted by resource utilizers (Fig. 3G). Succinate is linked to dysbiosis-related diseases and can be categorized as a primary metabolite crucial for the stability and resilience of gut microbiome ^38^, while carbon monoxide exhibits potent anti-inflammatory and immunomodulatory effects and can promote growth of beneficial bacteria^39^. Resource contributors were distinguished by the uptake of L-serine and the secretion of butyrate (Fig. 3G). Butyrate has been widely studied playing an essential role in intestinal homeostasis by increasing gut barrier integrity and modulating immune responses including low grade inflammation^40^. Previous studies have suggested that L-serine could provide a competitive fitness advantage for Enterobacteriaceae under inflammatory conditions^36,41^, which is consistent with our result showing that a large proportion of the members of the resource contributors came from Enterobacteriaceae. Moreover, L-lysine was preferentially consumed by resource contributors and active players but secreted by resource predators (Fig. 3G), and L-lysine is further known to promote immune tolerance in dendritic cells^42^. These results suggested that these four groups may have different ways of affecting gut health through metabolic exchange. Parallel analyses of metabolite competition yielded consistent clustering patterns (Supplementary Fig. 9A and B), confirming that these ecological groups exerted differential impacts on the gut ecosystem through both the consumption of limiting resources and the provision of bioactive metabolites.

### Influence of secondary metabolites on ecological differentiation

Since secondary metabolites play an essential role in bacterial fitness and ecological interactions, we conducted a targeted assessment of biosynthetic gene clusters (BGCs) in the four defined ecological groups based on their genome sequences. Our results showed that the distribution of BGCs was significantly associated with the division of these groups (Supplementary Fig. 10A). Specifically, NRP-metallophores were exclusively identified in resource contributors (Supplementary Fig. 10B). These molecules are known to bind essential metal ions like iron and zinc with high affinity^43^, suggesting that resource contributors could actively acquire resources from the environment, thereby enhancing their competitive fitness in nutrient-deprived niches. Conversely, cyclic-lactone-autoinducers were significantly enriched among active players (Supplementary Fig. 10B). As key signaling molecules in quorum sensing^44^, their prevalence suggests that active players may rely on coordinated chemical communication to orchestrate metabolic interactions and community dynamics. Remarkably, the compositional variance in BGCs (PCoA axes) exhibited strong congruence with the landscape of metabolite exchange and competition (Supplementary Fig. 11 and 12). This correlation implies a functional coupling between secondary metabolism (signaling and defense) and primary metabolism (nutrient exchange), suggesting that BGCs-encoded traits, such as chemical communication, may actively shape the metabolic interaction we observed.

### Application of the ecological group framework in inflammatory bowel disease (IBD)

To evaluate the resolution of our ecological group framework in capturing complex microbiome dynamics, we benchmarked it using longitudinal metagenomic data from the IBDMDB cohort, which is part of the Integrative Human Microbiome Project (HMP2)^45^. As the previous study reported^45^, the whole community composition tended to diverge more pronouncedly from baseline in both ulcerative colitis (UC) and Crohn’s disease (CD) compared to non-IBD (Fig. 4A). However, this broad trend masked different stability dynamics between disease subtypes and among ecological groups. Specifically, resource predators and resource contributors exhibited a more drastic temporal divergence from baseline compared with the whole community in both UC and CD (Fig. 4A and B). Intriguingly, resource utilizers in UC failed to exhibit the natural accumulation of dissimilarity observed in healthy individuals over time, suggesting a loss of normal temporal turnover (Fig. 4C).

**Fig. 4.**
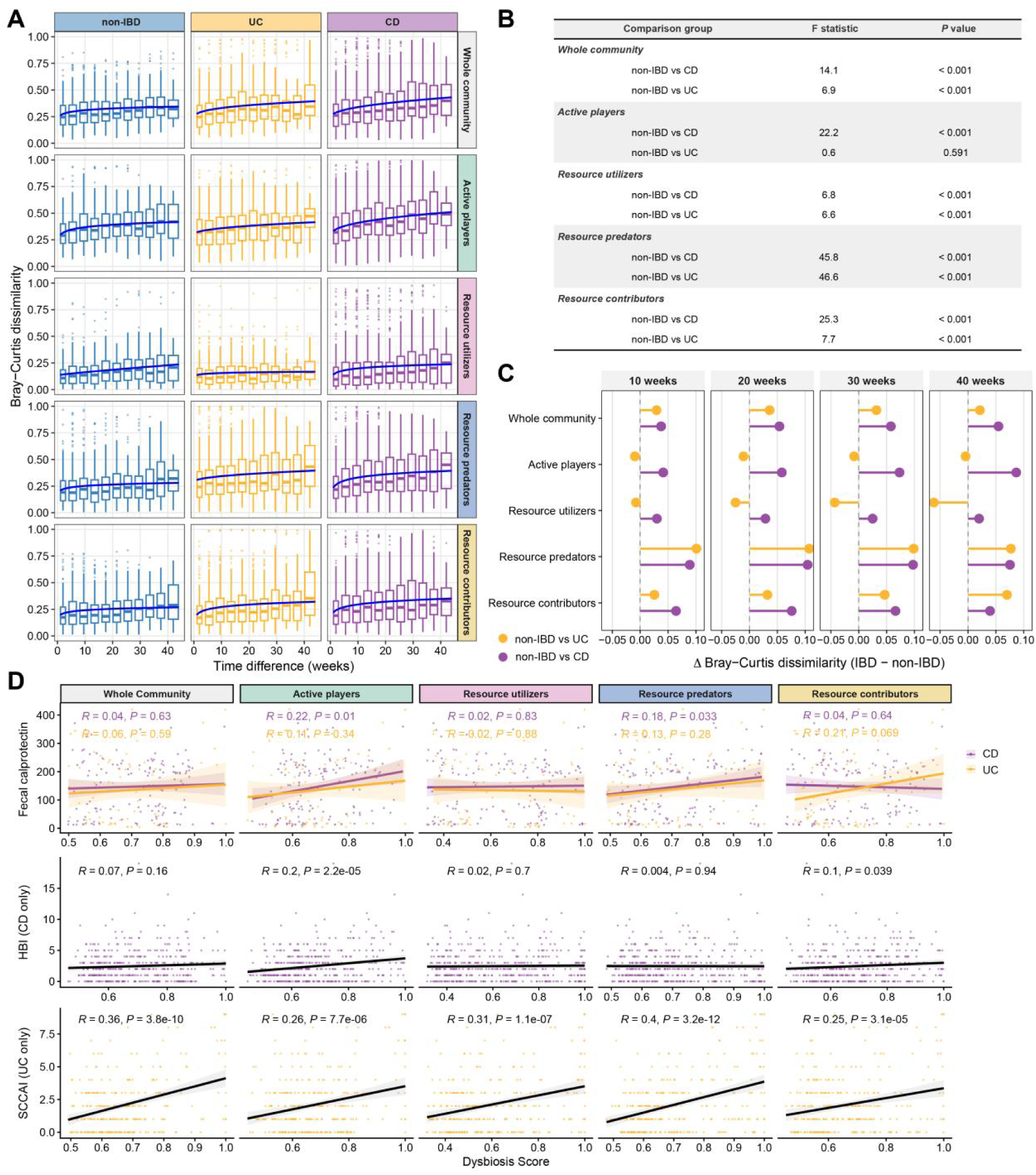
Longitudinal dynamics and clinical relevance of ecological groups in inflammatory bowel disease (IBD). **A** Temporal stability analysis showing Bray-Curtis dissimilarity between longitudinal sample pairs separated by increasing time intervals (lag in weeks). Trajectories represent the least-squares power-law fits. **B** Statistical comparison of the temporal turnover parameters between non-IBD and IBD subtypes. **C** Excess temporal instability in IBD cohorts relative to non-IBD controls, quantified as the difference in Bray-Curtis dissimilarity at 10, 20, 30, and 40-week time lags. **D** Associations between ecological groups-specific dysbiosis scores and clinical disease activity markers, including fecal calprotectin, Harvey Bradshaw Index (HBI) in CD patients, and Simple Clinical Colitis Activity Index (SCCAI) in UC patients.

We next evaluated whether these dynamic shifts reflected underlying changes in species co-occurrence patterns (Supplementary Fig. 13A). UC microbiomes were characterized by a universal synchrony, displaying significantly enhanced positive correlations both within and between all ecological groups compared with those in non-IBD. In contrast, significant shifts in species correlations were confined to the resource contributors in CD, which displayed weakened positive correlations internally and with other groups, indicating a breakdown in specific functional couplings. These results indicated contrasting microbial co-occurrence alterations between the two disease subtypes relative to non-IBD.

To assess the clinical utility of this framework, we calculated dysbiosis scores for each sample using the established algorithm based on both the whole community and ecological group-specific profiles^45^. We found that the group-based approach offered superior sensitivity, identifying a significantly higher proportion of dysbiotic samples, particularly within the active players (Supplementary Fig. 13B). Moreover, from a whole-community perspective, fecal calprotectin showed no significant correlation with dysbiosis scores in either UC or CD. However, partitioning of the microbiome revealed that dysbiosis within active players and resource predators was significantly and positively correlated with fecal calprotectin in CD (Fig. 4D). Similarly, while the Harvey-Bradshaw Index (HBI) in CD showed no association with whole-community dysbiosis, it correlated significantly with scores derived from active players and resource contributors (Fig. 4D). In UC, although the Simple Clinical Colitis Activity Index (SCCAI) correlated with both whole-community and group-specific scores, the resource predators yielded the strongest correlated coefficient (Fig. 4D). Collectively, these results confirmed that applying our ecological group framework to complex data provides a high-resolution lens for monitoring disease progression and stratifying patient phenotypes.

### High strain diversity at the level of metabolic interactions

Given the strain diversity characterizing the four ecological groups, we conducted an in-depth analysis of metabolic niches related to diversity. Among the 22 species (with ≥10 strains), we observed a clear differentiation in metabolic niches at the strain level. Specifically, different genomes within the same species exhibited distinct metabolic distance profiles concerning interacting with other microbial members (Fig. 5A). For instance, the strains in *Bifidobacterium longum* could be divided into two clusters, with Cluster 2 displaying the lower metabolic competition and higher complementarity with Cluster 1, but the opposite within Cluster 2 itself (Fig. 5B). Thus, our analyses revealed that Cluster 1 has a potential for secreting L-arabinose to support Cluster 2 (Fig. 5C). L-arabinose serves as a critical carbon source derived from dietary plant fibers, fueling the colonization and persistence of Bifidobacterium in the gut^46^. Furthermore, the metabolic exchange products of Cluster 1 and Cluster 2 with other species were largely non-redundant. For example, Cluster 1 has the potential to secret carbon monoxide and L-glutamate and take up butanoyl-CoA, ubiquinone-8, and cadmium, while Cluster 2 has the potential to take up D-glucose, ornithine, and L-homoserine (Fig. 5D). Additionally, the two clusters also exhibit obvious differences in competition with metabolites from other species (Supplementary Fig. 14). Consistent with our classification of ecological groups, Cluster 1 strains were all classified as resource utilizers, while the majority of Cluster 2 strains belonged to active players (Supplementary Fig. 15A). To validate the potential for the co-colonization of infraspecific lineages, we simulated and quantified growth rates under both mono-culture and pairwise co-culture conditions in a Western Diet environment (Supplementary Fig. 15B). Our results showed that Cluster 1 strains failed to grow in isolation, while their growth viability was fully restored when co-cultured with Cluster 2 strains. Conversely, pairings within Cluster 2 predominantly resulted in reduced growth rates, reflecting strong niche overlap and intense resource competition among conspecifics. Strikingly, the majority of Cluster 2 strains also exhibited enhanced growth rates when paired with Cluster 1 strains. This finding indicates a reciprocal mutualism, suggesting that these lineages may have evolved complementary metabolic niches to mitigate competition and facilitate robust infraspecific co-colonization^47^.

**Fig. 5.**
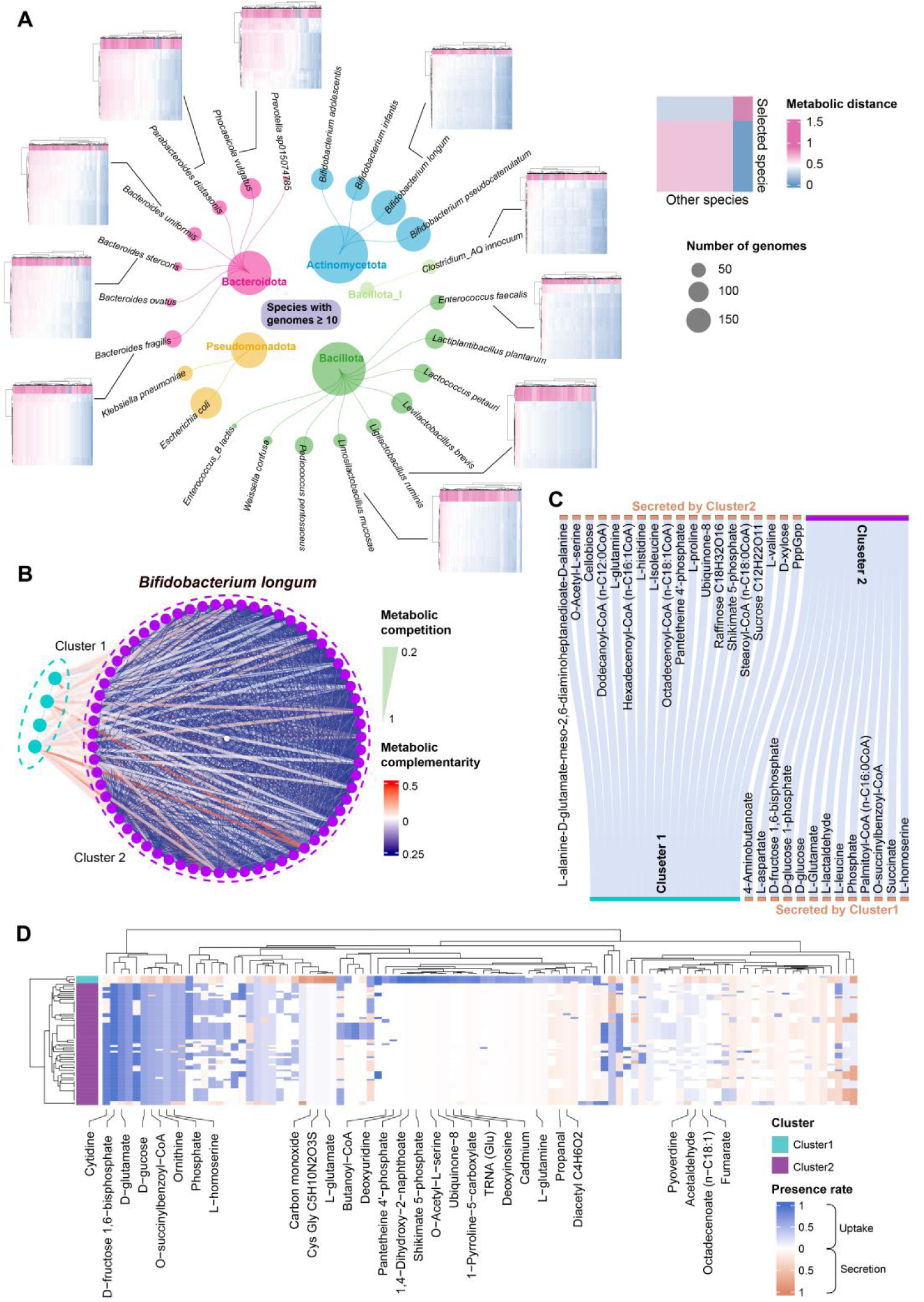
Strain diversity at the level of metabolic interactions. **A** The distribution of metabolic pattern differentiation within the genomes for species with genomes ≥10. **B** An example of metabolic pattern differentiation within *Bifidobacterium longum*. **C** Sankey plot showing the metabolite secretion and uptake between two genome clusters of *Bifidobacterium longum*. **D** Heatmap showing the metabolite secretion and uptake between two clusters of *Bifidobacterium longum* and other community members. The color scale indicates the potential, defined as the proportion of the total community with which a focal strain uptakes/secretes for a specific metabolite.

### Impacts of metabolic interactions on host health

We applied random matrix theory to construct co-occurrence networks based of three cohorts comprising healthy individuals, as well as metabolic networks based on complementarity and competition indices^34^. We found that the networks at the abundance level and those at the metabolic interaction level presented different patterns (Fig. 6A and Supplementary Table 9). We then identified keystones in each network using the Zi-Pi method, which is widely used in the field of microbial ecology^48,49^. The results showed that 28, 104, and 47 species belonged to keystone taxa in co-occurrence, metabolic complementarity, and competition networks, respectively. Among them, 22 keystone species were shared in metabolic competition and complementarity networks and were regarded as the keystones at the metabolic interaction level (Fig. 6B). Five species, including *Dorea_A longicatena*, *Faecalimonas phoceensis*, *Streptococcus hominis*, *Bifidobacterium infantis*, and *Phocaeicola plebeius_A*, were identified as keystone at both metabolic and abundance levels (Fig. 6C). Notably, these five species except *Streptococcus hominis* exhibited extremely high prevalences in healthy cohorts (Supplementary Table 10), indicating the universality of their key roles in stabilizing the gut microbiome. Furthermore, we developed gradient boosting and random forest classifiers to validate the importance of keystones in distinguishing diseased individuals from healthy individuals (Fig. 6D). We found that the classifiers constructed using keystones identified through the network at abundance level demonstrated moderate up to good diagnostic capabilities in certain diseases, including liver cirrhosis, ankylosing spondylitis, and atrial fibrillation, with area under the receiver operating characteristic curve (AUROC) between 0.8 and 0.9. After combining the keystones identified through the metabolic interaction network, the diagnostic abilities of the classifiers in all diseases even improved reaching AUROC above 0.9 for atrial fibrillation (Fig. 6D). These findings highlight that disease diagnostic models might benefit from incorporating ecological interaction networks at the metabolic level, which might capture functional dependencies that drive disease states.

**Fig. 6.**
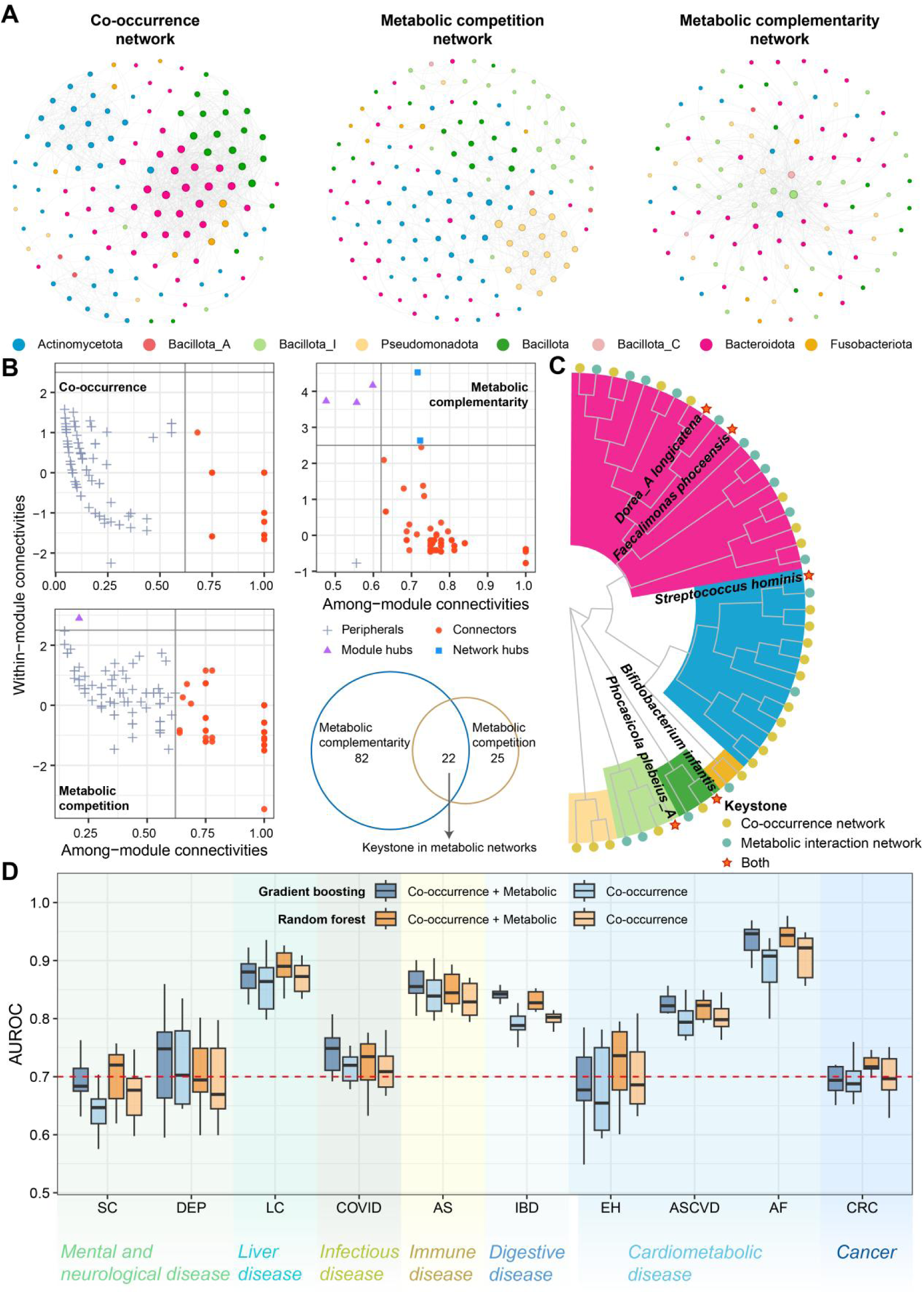
The integration of microbial co-occurrence and metabolic interaction networks. **A** Co-occurrence, metabolic competition, and complementary networks. **B** Identifications of keystone in co-occurrence and metabolic networks. **C** Taxonomic information of keystone. **D** Boxplots showing the performances of classifiers for disease diagnosis using keystone identified in only co-occurrence networks and in both co-occurrence and metabolic networks by random forest and gradient boosting, respectively. SC, schizophrenia; DEP, depression; LC, liver cirrhosis; COVID, COVID-19; AS, ankylosing spondylitis; IBD, inflammatory bowel disease; EH, essential hypertension; ASCVD, atherosclerotic cardiovascular disease; AF, atrial fibrillation; CRC, colorectal cancer.

## Discussion

Deciphering metabolic interactions is crucial for understanding microbiome assembly and function. By leveraging complete genomes to construct GEMs, we mapped the landscape of gut microbial interactions, revealing that genomic streamlining drives metabolic cross-feeding. We stratified the microbiome into four distinct ecological groups, including active players, resource predators, utilizers, and contributors, based on interaction asymmetries. This classification was mechanistically supported by distinct BGCs profiles and uncovered profound strain-level metabolic heterogeneity. Applying this framework to IBD cohorts revealed specific ecological instabilities and clinical associations obscured by whole community metrics. Furthermore, incorporating metabolic interaction keystones significantly improved diagnostic accuracy across multiple diseases. Our findings provide a high-resolution and mechanistic framework for decoding the functional architecture of the gut microbiome and its impact on host health.

Although the construction of GEMs from MAGs serves as a valuable alternative given that many bacteria remain unculturable in the laboratory, and their credibility has been successfully validated^50^, the quality of MAGs, particularly their completeness, could significantly impact the accuracy of GEMs^51^. A previous study suggested that incomplete metabolic networks could introduce biases in flux predictions, as blocking even a few reactions significantly altered downstream pathways^52^,which indicates that constructing GEMs with complete genomes is essential. Our study revealed that GEMs constructed using complete genomes enable the recovery of a substantial number of metabolic reactions, predominantly related to transport. This could have a significant impact on metabolite flux, which in turn influences metabolite competition and exchange. Considering that metabolic modeling has been used to distinguish patients from controls based on individual-specific metabolic fluxes in common diseases^24^, complete genomes are also expected to significantly enhance this discriminatory performance.

While habitat filtering drives the co-occurrence of species with similar metabolic preferences^36,53^, their metabolic interdependence is governed by the "Black Queen Hypothesis"^54^. This posits that bacterial genomes may undergo reductive evolution to eliminate non-conserved genes, thereby improving growth efficiency and reducing metabolic costs, while such genomic streamlining often creates metabolic dependencies that must be compensated through increased cross-feeding interactions with other community members^55,56^. As expected, we observe a clear genomic gradient where strains possessing larger genomes retain costly biosynthetic pathways to support metabolically streamlined counterparts, leading to widespread specialization and monopolization of scarce resources^57^. Crucially, the landscape of BGCs strikingly mirrors the patterns of primary metabolic exchange, suggesting that primary nutrient exchange and secondary chemical signaling are not isolated processes but coupled with each other to ensure microbially ecological stability^58,59^.

The clinical utility of this metabolically defined framework is highlighted when applied to complex disease states like IBD. While traditional taxonomic profiling often blunts the distinction between disease phenotypes, our ecological group-based approach acts as a high-resolution lens, uncovering specific differences in UC and CD that were previously masked^45^. The finding that dysbiosis scores derived from specific groups correlated more strongly with clinical biomarkers than those from whole community suggested that disease progression could be driven by the breakdown of specific functional modules rather than generalized community shifts^60^. Moreover, this resolution extended beyond the species level to the strain level. The reciprocal mutualism we identified between *B. longum* clusters demonstrated that the species is often too coarse an ecological unit^61^. Such intraspecific metabolic divergence likely evolved to mitigate competition, allowing lineages to co-colonize the host by partitioning metabolic niches, a nuance that is critical for the design of precision probiotics^62,63^.

To address the disconnection observed between co-occurrence patterns and metabolic potential^34,64^, we integrated metabolic competition and complementarity networks with empirical abundance data. This approach moves beyond simple correlation^65-67^, which can be confounded by environmental selection^68^, to identify keystone taxa based on network structure rather than mere abundance^69^. The biological validity is exemplified by *B. infantis*, which emerged as a dual keystone and its topological importance aligns perfectly with its established capacity to suppress inflammation and resist pathogen colonization^70^. Importantly, unlike other studies that use biomarkers, we leveraged network keystones to construct disease classifiers. This reflects that the stability of gut microbial communities is critical for host health, and keystone taxa play a pivotal role in maintaining stability^71-73^. Disease classifiers incorporating metabolic keystones significantly outperformed traditional models, highlighting the critical role of gut microbially metabolic interaction in host health^74^.

In summary, our study bridges the gap between metabolic potential and ecological reality. By establishing the necessity of complete genomes, exploring the evolutionary rules of cross-feeding, and validating the clinical power of ecological groups, we provide a mechanistic blueprint for understanding gut microbiome assembly. These insights may pave the way for next-generation therapeutics, where interventions are designed not just to introduce missing taxa, but to repair the fractured metabolic networks that underpin host physiology. Furthermore, as long-read sequencing facilitates the recovery of complete MAGs^75^, our approach provides a feasible solution for decoding the metabolic interactions of complex communities that exceed the culture-dependent limits. However, there were several limitations in our study. First, the reliance on strains isolated from healthy individuals may limit our ability to capture dysbiosis-specific metabolic traits or strain-level adaptations driven by disease pressures. Expanding the culture collection to include patient-derived isolates will be crucial for future disease modeling. Second, the *in silico* interactions in our results await causal validation. Future studies should leverage *in vitro* co-culture assays and gnotobiotic models to experimentally verify the impact of metabolic interactions and identified keystones on host physiology.

## Methods

### Sample collection and preservation

In this study, we recruited 74 healthy volunteers who had no history of intestinal diseases for fecal sampling. The recruited individuals had not been treated with any antibiotics in six months prior to sampling and not been taking any other drugs during the last month before sampling of fecal material. The sampling was approved by the Institutional Review Board on Bioethics and Biosafety of BGI (Approval No. BGI-IRB17005-T1). All experimental protocols comply with the Declaration of Helsinki, and informed consent has been obtained from all participants. Fecal samples were collected in 2 ml of a protective solution containing 25% anaerobic glycerol (diluted with PBS) and 0.5 g/L sodium sulfide (Na2S), 0.25 g/L cysteine, and 1 ml/L resazurin liquid to ensure its activity, and then immediately transported on dry ice to a freezer at -80 ℃ for storage. Information regarding the strain culture conditions, sources, genomic information, and species annotation can be found in Supplementary Tables 1-5.

### Bacterial isolation and culture

In our previous studies, we found that a few culture media were sufficient to obtain the vast majority of species^76,77^. Therefore, in this study, we used only Modified Peptone Yeast Glucose (MPYG), Brain Heart Infusion (BHI), Gifu Anaerobic Medium (GAM), de Man, Rogosa, and Sharpe (MRS) medium, and Weizmannia Count Agar (WCA) for isolation and culture. Samples were immediately transferred to an anaerobic incubator (Bactron IV-2, Shellab, USA), homogenized in pre-reduced phosphate-buffered saline (PBS, supplemented with 0.1% cysteine), then diluted and spread onto agar plates containing different media.

Plates were incubated at 37 ℃, 90% N_2_, 5% CO_2_, and 5% H_2_ for 2-3 days under anaerobic conditions. Single colonies were picked and streaked onto new plates to obtain monoclonal strains. All strains were preserved in a 20% (v/v) glycerol suspension containing 0.1% cysteinate at -80°C. The detailed specific growth conditions (temperature, gas composition, incubation time) are provided in Supplementary Table S1.

### Genome sequencing, assembly, quality assessment, and gene prediction

To account for the diversity of strains, we selected 2-3 representative strains of the same species from each fecal sample for short-reads (DNBSEQ-T7) and long-reads (CycloneSEQ-G100) sequencing. In our study, we isolated bacterial strains during the experiment and performed short-read and long-read sequencing on each batch of bacterial culture. Short-reads sequencing was performed as previously described^78^. For the long-reads sequencing, the DNA from each strain was extracted using the Magen MagPure DNA Kit for high-throughput applications. Library preparation and sequencing with CycloneSEQ were conducted following the manufacturer’s guidelines and protocols. Libraries were quantified using a Qubit fluorometer and sequenced on the CycloneSEQ-G100 platform according to standard sequencing protocols. Long-read data were filtered using NanoFilt (v2.8.0) with parameters “-q 10 -l 1000” to retain reads longer than 1,000 bp with a quality score ≥ Q10. Short-read data were processed using Fastp (v1.0.1) with default parameters, except for a length cutoff set to 50 bp^79,80^. Short-read assembly was performed using Unicycler (v0.5.1), with only the short read sequences "-1" and "-2" as input, and "--depth_filter" set to 0.01 to remove low-depth contigs. For hybrid assembly, "--depth_filter" was also set to 0.01, and the long read sequence "-l" was added as input, with all other parameters set to their default values.

The circularity of each genome was evaluated using the assembly results from Unicycler and Flye^81,82^. Genome quality was assessed using CheckM2^83^, circular genomes with >95% completeness and <5% contamination were retained for further analysis. The proportions of BGCs were retrieved using the antiSMASH^48^.

### Phylogenetic and taxonomic analyses

All genomes were taxonomically annotated using GTDB-Tk (v2.4.0) ‘classify_wf’ function against the GTDB database r220. The genome maximum-likelihood phylogenetic tree was constructed using ‘infer’ function of GTDB-Tk based on 120 bacterial marker genes and visualized by iTOL. The phylogenetic distance was calculated using the cophenetic function in R package ape.

### Species prevalence and abundance

Human gut metagenome sequencing data of healthy individuals from Chinese (a part of 4D-SZ), the Netherlands, and the HMP cohorts were downloaded. Furthermore, we collected human gut metagenome sequencing data of 10 disease cohorts for disease diagnostic analysis, including schizophrenia, depression, liver cirrhosis, COVID-19, ankylosing spondylitis, inflammatory bowel disease, essential hypertension, atherosclerotic cardiovascular disease, atrial fibrillation, and colorectal cancer. Detailed source information is placed in Data availability. We used dRep (v3.4.2) to construct species-level clusters of our complete genomes with option "--ignoreGenomeQuality -sa 0.95". After that, we used these 199 species-level clusters to build a custom genome database by Kraken (v2.1.257) and Bracken (v2.558). For these cohort data, we first used fastp (v1.0.1) to remove low-quality sequences with option "--trim_poly_g, --poly_g_min_len 10, --trim_poly_x, --poly_x_min_len 10, --cut_front, --cut_tail, --cut_window_size 4, --cut_mean_quality 20, --qualified_quality_phred 15, --low_complexity_filter, --complexity_threshold 30, --length_required 50", and then use bowtie2 (v1.3.1) to remove host sequences (GRCh38) with option "--very-sensitive". The 199 species-level clusters were built as a custom genome database by Kraken v2.1.257 and Bracken v2.558. To calculate prevalence, a threshold of 0.01% relative abundance was used to define the occurrence of a cluster in one sample^84^.

### GEMs construction

GEMs for all genomes were constructed using CarveMe (v1.5.1) with default parameters^27^. In contrast to conventional bottom-up methods that necessitate well-defined growth media, manual curation, and gap-filling processes, the top-down approach employed by CarveMe eliminates reactions and metabolites deemed absent in the manually curated universal template^85^. The input for CarveMe was coding sequences of each genome, which was predicted based on Prokka^86^. The interactions between pairwise genome-scale metabolic models were predicted using PhyloMint^85^. All modeling software mentioned above required the installation of an optimization solver, which for our study is IBM CPLEX Optimizer (ILOG COS 20.10 Linux x86-64 version). Central to this analysis is the concept of seeds or seed sets. Biologically, the seed set represents the specific profile of essential nutrients and metabolites that a microorganism is unable to synthesize internally and must therefore acquire from its external environment to synthesize its cellular biomass. Computationally, these seeds are identified based on the graph topology of the metabolic network. They are defined as the minimal set of exogenous substrates required to activate the metabolic graph, often determined by analyzing strongly connected components where specific metabolites act as source nodes^85^ (PMID: 33125363). Based on these identified seed sets, metabolic overlap was measured to calculate the metabolic competition index. When considering two genomes A and B, the fraction of genome A’s seed set that overlaps with genome B’s seed set is normalized by the weighted sum of confidence scores for A’s seed set. For the metabolic complementarity index, the fraction of genome A’s seed set found in genome B’s metabolic network but not in genome B’s seed set is normalized by the total size of genome A’s seed set in B’s network, indicating the potential for genome A to utilize genome B’s metabolic outputs. The differences of metabolic competition and complementarity indices between complete and draft genomes were tested by Kruskal-Wallis tests. Both indices are asymmetric and computed using seed sets from genome-scale metabolic models^85^. Given the negative correlation observed between competition and complementarity, we calculated a combined metabolic distance score using the formula: Metabolic distance = 1 - (Competition index-(Complementarity index)). This metric integrates both interactions to reflect metabolic niche preference: a smaller distance indicates significant metabolic niche overlap and similar resource preferences, whereas a larger distance reflects niche differentiation^36^. Linear regression models were applied to test the effects of genome size and phylogenetic distance on metabolic interaction indices. To assess the specific impact of functional gene categories (COG) on metabolic interactions, we used linear mixed-effects models following the previously established methodology^34^. The goal was to estimate the effects of biological functions on metabolic potential while controlling for genomic structural factors. Prior to analysis, all continuous variables were standardized (scaled) to a mean of 0 and a standard deviation of 1. The pairwise differences in COG functional categories were treated as fixed effects. The pairwise genome size difference and phylogenetic distance were modeled as random effects. This design allowed for the estimation of functional gene effects while accounting for the baseline variation introduced by genome size and phylogenetic divergence. All models were fitted using the lme4 package in R, and assumptions of normality and homoscedasticity were verified.

### Division of ecological groups based on metabolic interactions

To comprehensively delineate the ecological roles of strains, we developed a comparative framework based on the asymmetry of metabolic interactions. Specifically, for each focal strain, we aggregated metabolic interaction indices into two distinct distributions, including those derived from interactions where the focal genome acted as the initiator (active role) and those where it acted as the partner (passive role). We assessed the statistical significance of the divergence between these two distributions using the Wilcoxon test, employing a stringent threshold of *P* < 0.001 to minimize false discoveries. To further ensure the robustness of our classification, we performed a bootstrap resampling analysis (n = 100 iterations), classifying a strain only if its group assignment remained consistent across iterations. Based on the statistical sign and significance of differences in metabolic competition and complementarity indices, strains were categorized into four distinct ecological groups: 1) active players, which showed significantly higher active competition and complementarity, indicating a proactive strategy in both resource acquisition and metabolic exchange; 2) resource predators, which showed higher active competition but lower active complementarity, suggesting dominance in competitive resource seizure while contributing minimally to cooperative exchange; 3) resource utilizers, which displayed higher active complementarity but lower active competition, reflecting a high capacity for initiating metabolic complementarity while avoiding competitive exclusion; 4) resource contributors, which had significantly lower active metrics in both, suggesting that they could primarily function as passive resource hubs or recipients of competition, driving community turnover through their susceptibility. This robust computational framework enabled a high-resolution classification of genomes based on their metabolic interaction profiles, facilitating the identification of distinct ecological roles within the gut microbiome.

### Community knockout analysis

To quantify the structural contribution of ecological groups, we performed computational knockout experiments on randomized synthetic communities. Metabolic interaction potentials were mapped for all 1,150 GEMs based on the directionality of exchange reactions. Synthetic communities (n = 20) were assembled by random sampling of 100 strains, constrained to include representatives from all four ecological groups. We defined a metabolic support score (MSS) to quantify the fraction of a strain’s auxotrophies met by the community environment. Specifically, for a community (*C*), the available metabolic pool (*P_comm_*) was defined as the union of a basal medium and the secretion profiles of all members 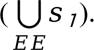 The MSS was calculated as 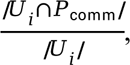 where *U_i_* represents the uptake potential of strain $i$. Stability loss was quantified as the relative reduction in MSS following the targeted removal of all strains belonging to a specific group 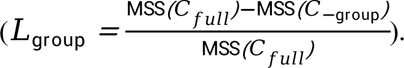

### Determination of potentially exchangeable metabolites

Using the PhyloMint algorithm, seed metabolites were defined by strongly linked components and represent exogenous substrates obtained^85^. For example, when considering two genomes A and B, potentially exchangeable metabolites were defined as those that were found in the seed set of genome A and present in the metabolic network of genome B, but not in the seed set of genome B. This indicates that B can synthesize these metabolites, while A requires them as exogenous substrates. The metabolite profiles were taken directly from the corresponding GEMs generated by CarveMe with the BiGG Models database. PCoA and permutational multivariate analysis of variance were used to test the differences of the metabolite exchange/competition between genomes in four ecological groups and those from other species among four groups using the vegan R package.

### Ecological group framework in IBD

To benchmark the utility of the defined ecological groups against traditional whole community metrics, we analyzed longitudinal shotgun metagenomic profiles from the IBDMDB cohort of HMP2, comprising temporally resolved samples from non-IBD, UC, and CD subjects. To characterize the temporal stability of the gut microbiome, we calculated the Bray-Curtis dissimilarity between all longitudinal sample pairs from the same subject. Sample pairs were stratified by the time interval (lag) separating them, ranging from 0 to 45 weeks. We applied a non-linear regression model to quantify the divergence of community composition over time for the whole community and each ecological group by fitting a power-law curve with free intercept by least-squares. The temporal stability of IBD subtypes (UC and CD) was compared against the non-IBD baseline using F-tests on the fitted model parameters. To explicitly quantify the loss of stability in disease, we calculated the excess instability ( Δ dissimilarity) by subtracting the mean fitted dissimilarity of the non-IBD from that of the IBD at discrete time points (10, 20, 30, and 40 weeks).

To investigate potential shifts in microbial co-occurrence patterns under disease conditions, we analyzed the distributions of Spearman’s correlations of pairwise species. *P*-values were adjusted for multiple testing using the Benjamini-Hochberg (FDR) procedure, and only statistically significant correlations (FDR < 0.05) were retained for analysis. The strength of these associations (*R*) was then quantitatively compared across disease states and ecological groups using the Kruskal-Wallis test.

We adapted the dysbiosis score algorithm previously described^45^. Briefly, a reference set was defined using samples from non-IBD subjects (collected after week 20 to ensure stability). For each query sample, the Bray-Curtis dissimilarity was calculated against all samples in the reference set (excluding samples from the same subject to prevent self-matching bias). The dysbiosis score was defined as the median of these dissimilarity values. To determine the prevalence of dysbiosis, we established a threshold for binary classification in which a sample was designated as dysbiotic if its dysbiosis score exceeded the 90th percentile of the scores observed in the non-IBD reference population. This procedure was performed independently for the whole community and for each of the four defined ecological groups, generating five distinct scores per sample. Associations between these group-specific dysbiosis scores and clinical metadata, including fecal calprotectin, HBI for CD, and SCCAI for UC, were assessed using linear regression models.

### Cultivating simulation

To investigate the coexistence potential among distinct *B. longum* lineages, we simulated growth in both mono-culture and pairwise co-culture under Western Diet conditions using the MICOM (v0.32.2)^23^. Cooperative metabolic interactions were modeled using the cooperative_tradeoff method (fraction = 0.5) to prevent competitive exclusion artifacts. Metabolic interactions were quantified as the relative change in growth rate as (Growth_co-culture - Growth_mono-culture) / Growth_mono-culture. Notably, rescue events (transition from zero monoculture growth to viability) were imputed with a fixed maximum positive score (0.5) to resolve mathematical indefinability.

### Metabolic interaction and co-occurrence network constructions

Since our goal was to construct undirected metabolic interaction networks, we symmetrized metabolic complementarity and competition indices for each genome pair. Specifically, we retained the maximum value of the two directional measurements to represent the strongest potential for metabolic competition or complementarity between the pair. These symmetrized values served as the adjacency matrices for the subsequent construction of undirected metabolic interaction networks^34^. We then determined the cutoff of these adjacency matrices based on the random matrix theory (RMT)-based approach using the RMT cutoff tool on iNAP website^51^. Metabolic complementarity and competition networks were visualized using the Gephi platform.

Regarding the co-occurrence network, species abundance profiles were generated using Kraken v2.1.2 and Bracken v2.5 as described in the section of Species prevalence and abundance. We normalized the absolute abundance table and obtained the relative abundance table. Using this table, the paired Spearman’s correlation of every two species was calculated. Based on this, the co-occurrence network was constructed using the RMT-based approach. Before applying the RMT cut-off tool, we removed species that were absent (zero abundance) in more than half of the samples using the majority selection tool on the iNAP website^51^. This step serves to eliminate the deviations caused by the influence of too many zeros from the relevant calculations. The separated noises form a network representing genomic pairs with high abundance correlations, which inferred their co-occurrence. The network was also visualized using the Gephi platform.

### Identification of putative keystone

To identify topologically important species, we performed the same topological role analysis on the co-occurrence, metabolic complementarity, and metabolic competition networks. The nodes in each network were classified into four topological roles based on their within-module connectivity (Zi) and among-module connectivity (Pi) values: module hubs (Zi ≥ 2.5, Pi < 0.62), network hubs (Zi ≥ 2.5, Pi ≥ 0.62), connectors (Zi < 2.5, Pi ≥ 0.62) and peripherals (Zi < 2.5, Pi < 0.62). Except for peripherals, the other three categories were regarded as putative keystone due to their crucial roles in stabilizing the network structure^49,87^. These putative keystone taxa identified from different networks were subsequently used as features for the machine learning classifiers.

### Machine learning

We employed random forest and gradient boosting models to classify case and control from 10 types of disease cohorts, with keystone (identified in only co-occurrence networks and in both co-occurrence and metabolic networks) serving as features, using the R package randomForest, gbm, and caret. Model training and hyperparameter tuning were performed on 70% of the data using a 5-fold cross-validation, while the other 30% were used for testing with the best hyperparameter setting. The whole procedure was then repeated 10 times with independent seeds. Model performance was evaluated using the AUROC.

## Supporting information

SFigures

## Data availability

Our complete bacterial genomes and high-quality plasmid and phage genomes have been uploaded at CNSA (CNP0007680). The the public cohorts we used in this study included ankylosing spondylitis (ERP111669), schizophrenia (ERP111403), depression (SRP337805), essential hypertension (ERP023883), liver cirrhosis (ERP005860), COVID-19 (DRP009646), atrial fibrillation (ERP110580), colorectal cancer (ERP012177, DRP004793, SRP320766), atherosclerotic cardiovascular disease (ERP023788), Chinese cohort (a part of 4D-SZ) (https://db.cngb.org/search/project/CNP0000426/), HMP (including inflammatory bowel disease) (https://portal.hmpdacc.org/), the Netherlands cohort (https://ega-archive.org/studies/EGAS00001005027).

## Author contributions

L. X., Y.Zou. and F.Z.supervised and led the project. Y.G. initiated the idea and conceived the study. Y.G., H.W., designed and executed the analysis. H.L., M.W. and Z.W. download the cohort data. J.Y., Y.Zhong. and H.Z. preparation and cultivation of strains. T.Z., Y.D. and Y.Zhang. constructed the methods of assembly. L.Y., Y.X., J.Z., Y.W., J.W., X.R., Y.W., X.S., Y.T., S.C., B.W., X.J., C.L., X.X., J.X. Y.M and P.G. collected information and interpreted data. J.Y., F.L., Z.Y., and W.L. constructed bacterial sequencing library. Y.G. H.W., J.Y. and T.Z. wrote the manuscript. Y.Zou., L.X. and Y.S. reviewed and revised the manuscript. All authors reviewed and approved the final version of the paper.

## Supplementary figures

**Supplementary Fig. 1.**
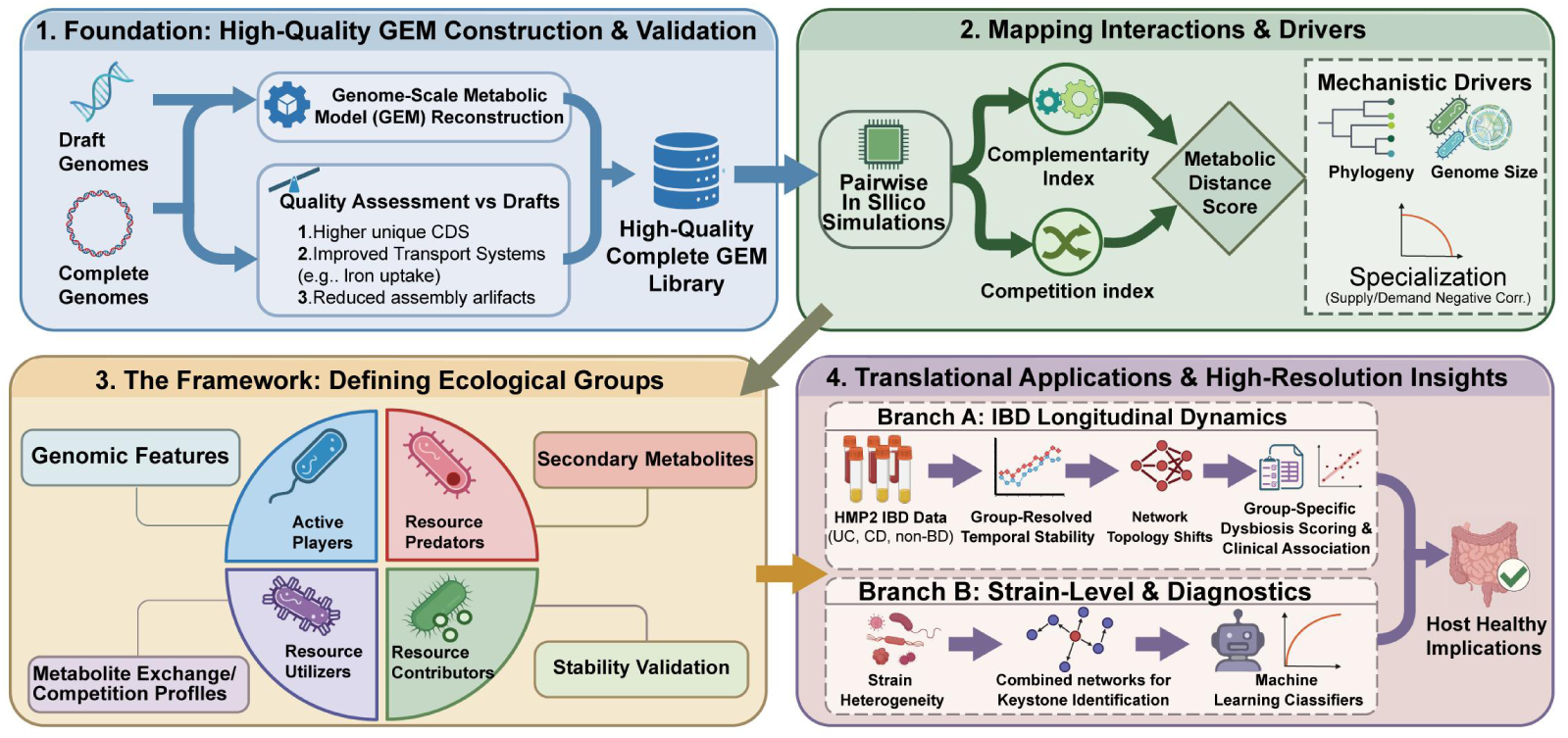
Exploring metabolic interaction patterns and their ecological effects. Workflow developed for metabolic interaction analysis using complete genomes.

**Supplementary Fig. 2.**
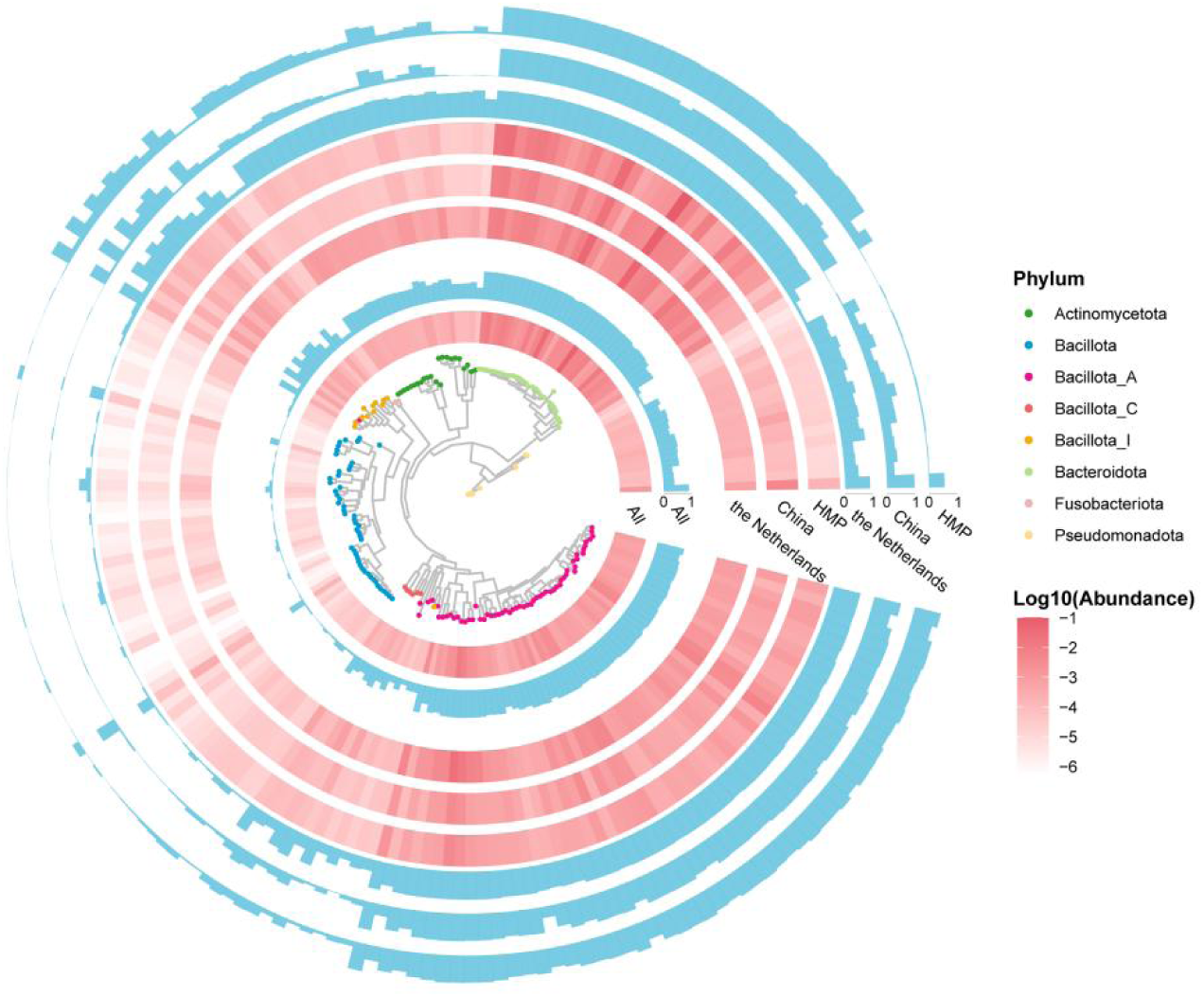
Distributions of species in healthy individuals. Abundance and prevalence of species in cross sectional cohorts of healthy individuals in China, HMP, and the Netherlands. The depth of the red color represents the abundance level, and the height of the blue column represents the prevalence level.

**Supplementary Fig. 3.**
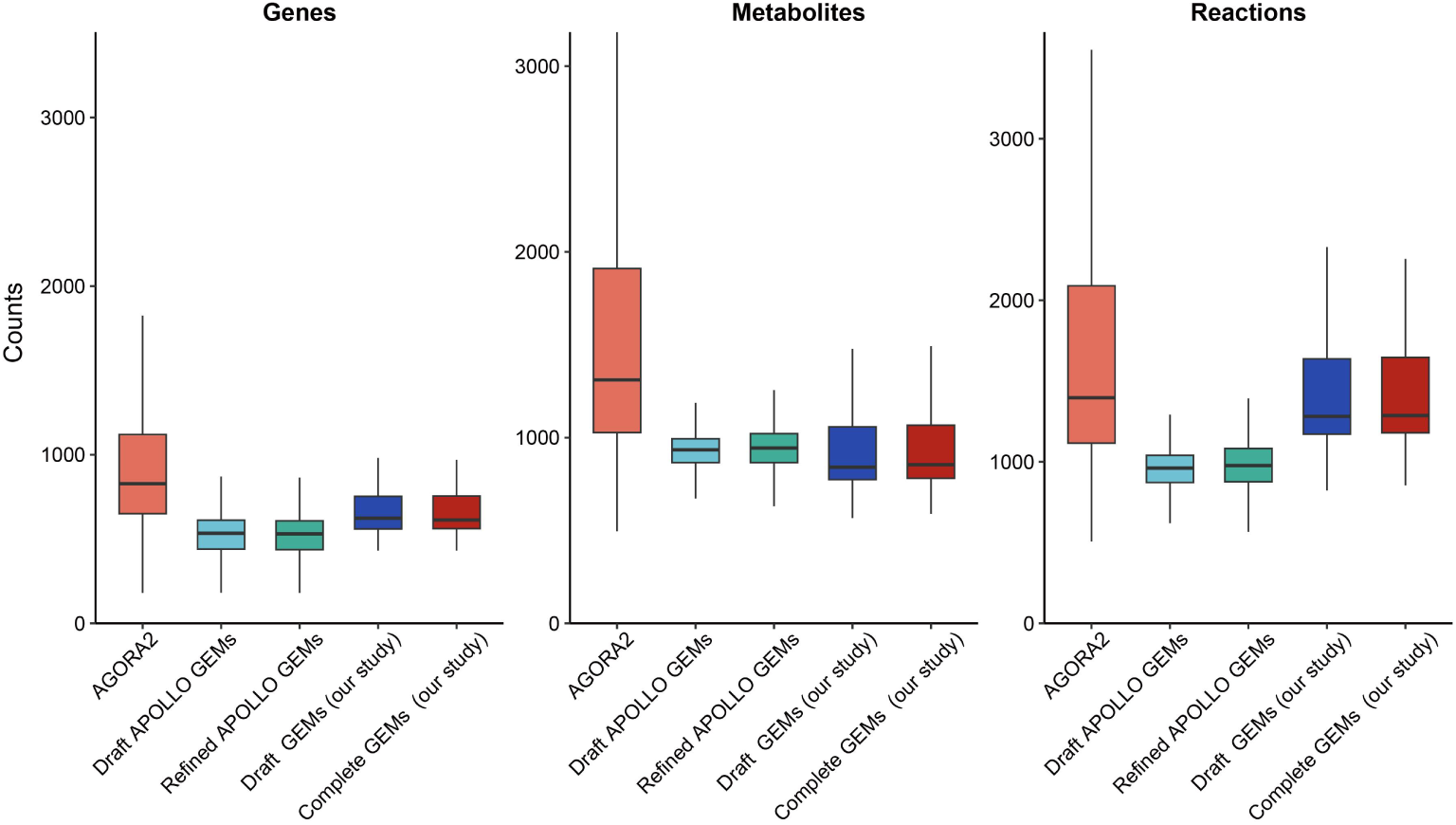
Comparison of different GEMs databases. The differences in the number of genes, metabolites, and reactions among different GEMs databases.

**Supplementary Fig. 4.**
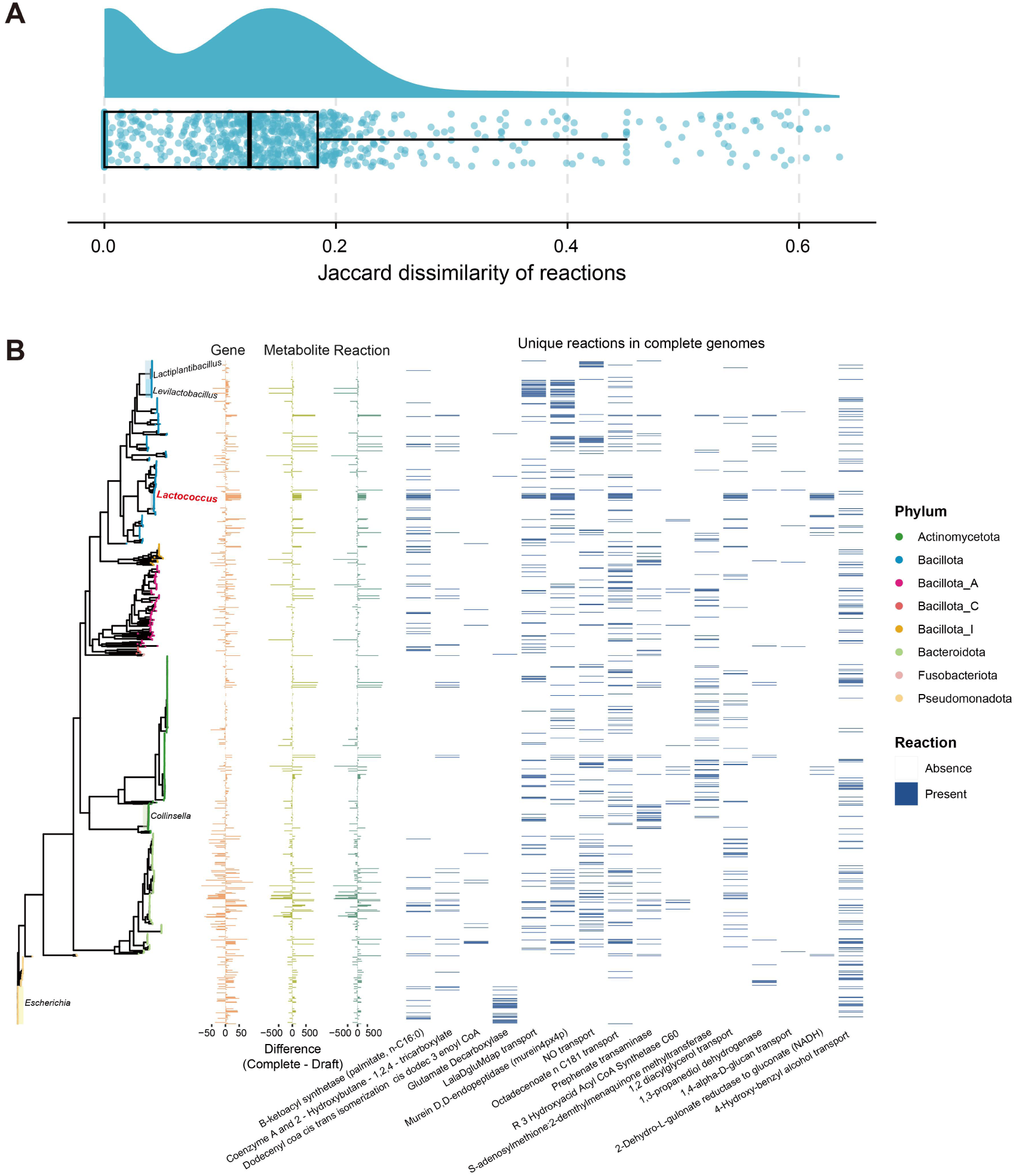
Differences between complete and draft genome-based GEMs. **A** Jaccard dissimilarity of reactions between complete and draft genome-based GEMs for each genome. **B** Variations between complete and draft genome-based GEMs. Phylogenetic tree colored by bacterial phylum is shown on the left, bar plots in the middle show the differences in the counts of genes, metabolites, and reactions for each strain, and heatmap on the right shows the unique reactions with high occurrence rates discovered in complete genome-based GEMs compared with these from draft genomes.

**Supplementary Fig. 5.**
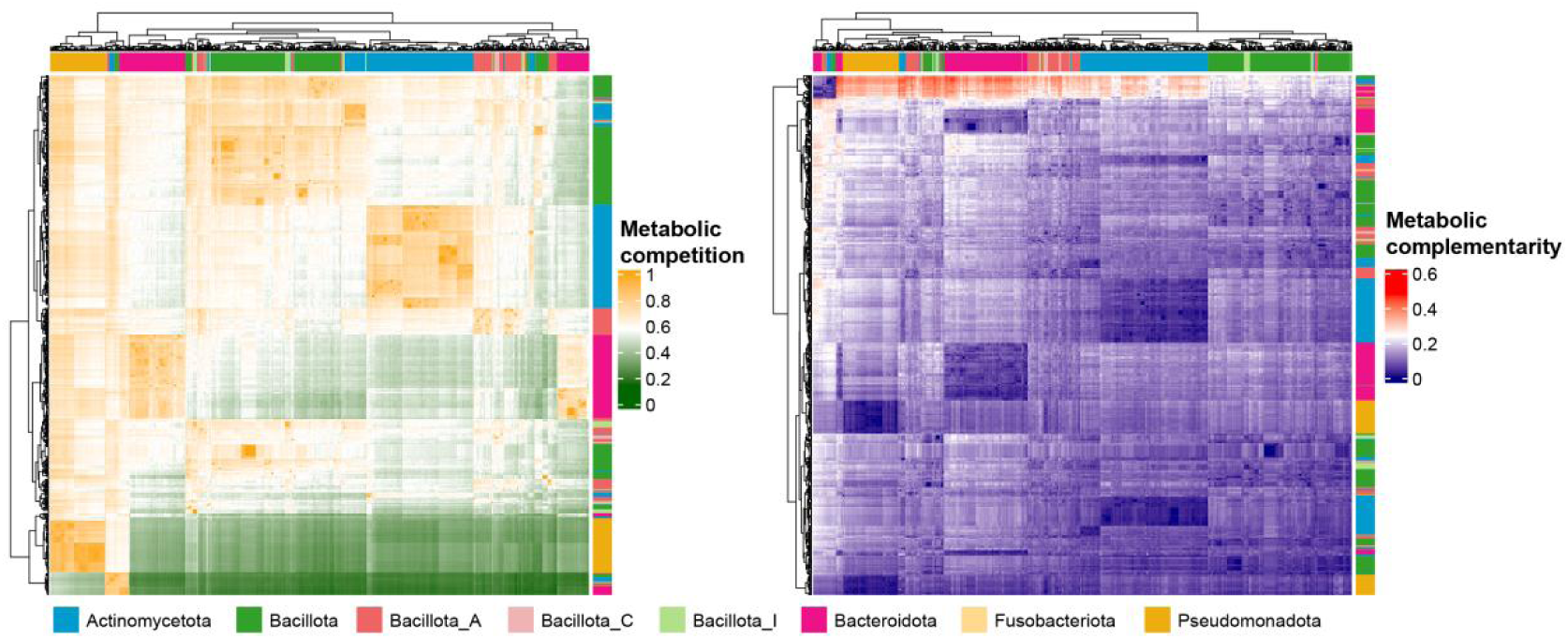
Metabolic interaction patterns. **Heatmaps showing t**he metabolic competition (**A**) and metabolic complementarity indices between every pair of genomes (**B**).

**Supplementary Fig. 6.**
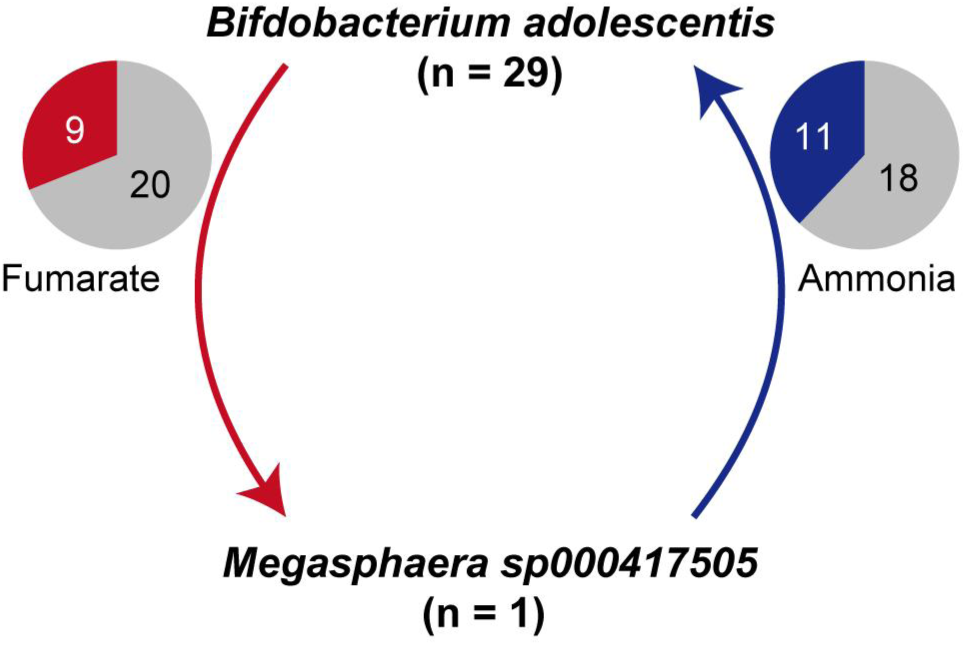
An example of metabolite exchange. Schematic diagram showing metabolic exchange of fumarate and ammonia between *Bifdobacterium adolescentis* and *Megasphaera sp000417505.* Pie plots show how many genes in *Bifdobacterium adolescentis* could be involved in these exchanges.

**Supplementary Fig. 7.**
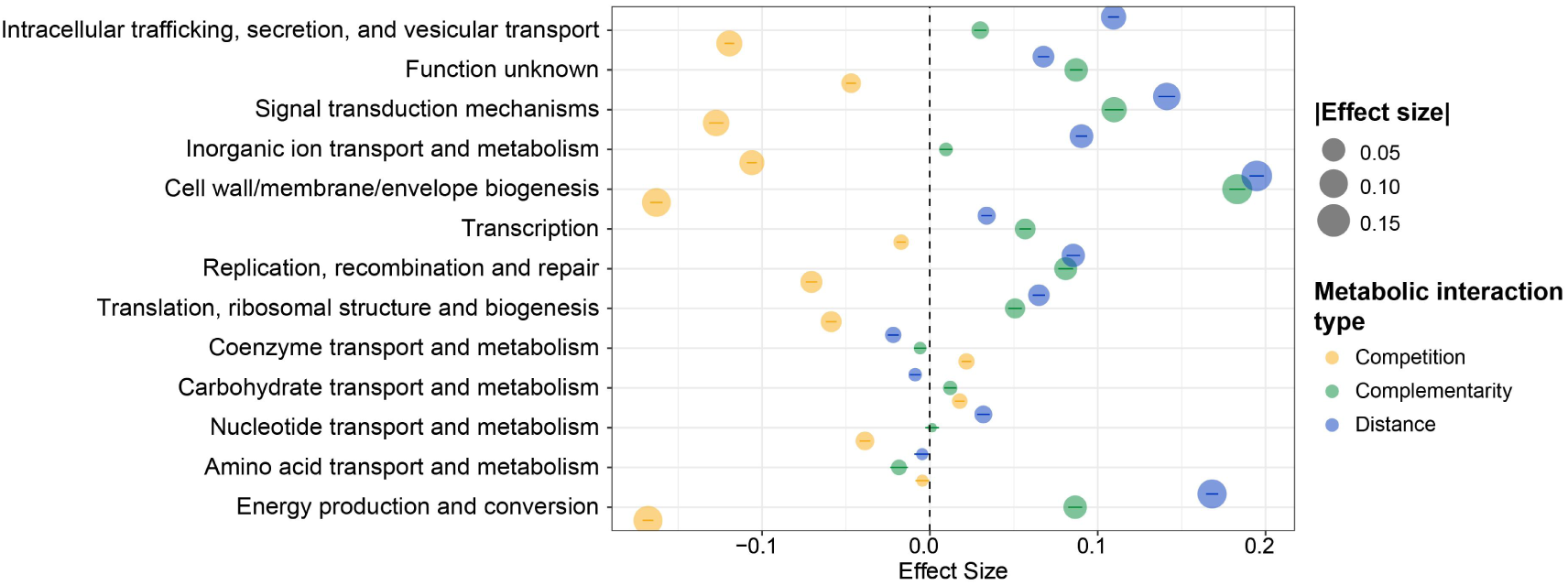
Impacts of bacterial function on metabolic interaction. Effect of COG biological function proportion on metabolic competition, complementarity, and distance using linear mixed-effects models. Bar length and error bar indicate the mean values and standard errors of the estimated effect sizes.

**Supplementary Fig. 8.**
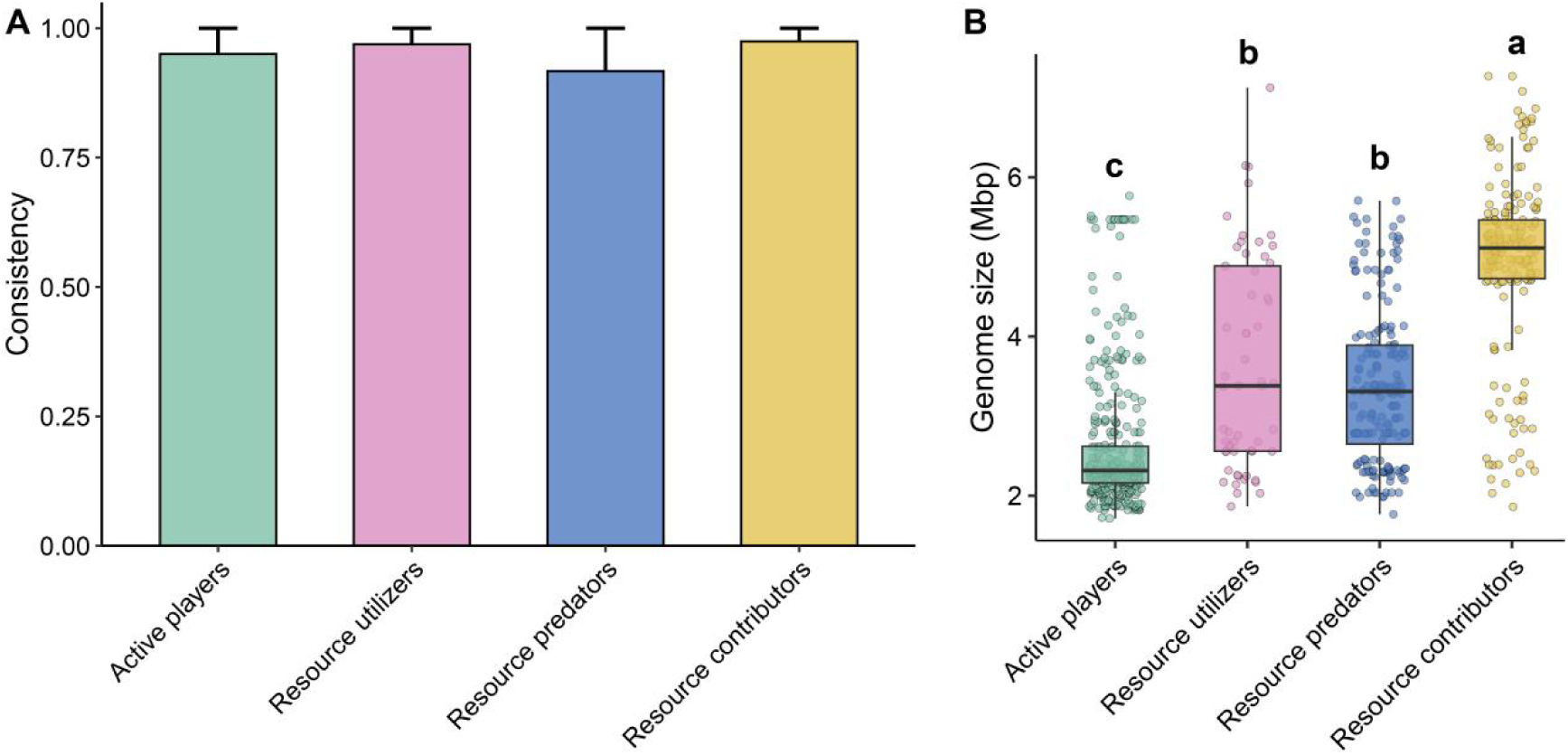
Robustness validation of ecological group assignments and genome sizes of groups. **A** To assess the stability of the inferred ecological roles (Active players, Resource utilizers, Resource predators, and Resource contributors), a bootstrap analysis was performed with 100 iterations. The skewness towards high consistency scores confirms that the classification of ecological groups is statistically robust. **A** Boxplots showing the genome sizes of strains across the four groups.

**Supplementary Fig. 9.**
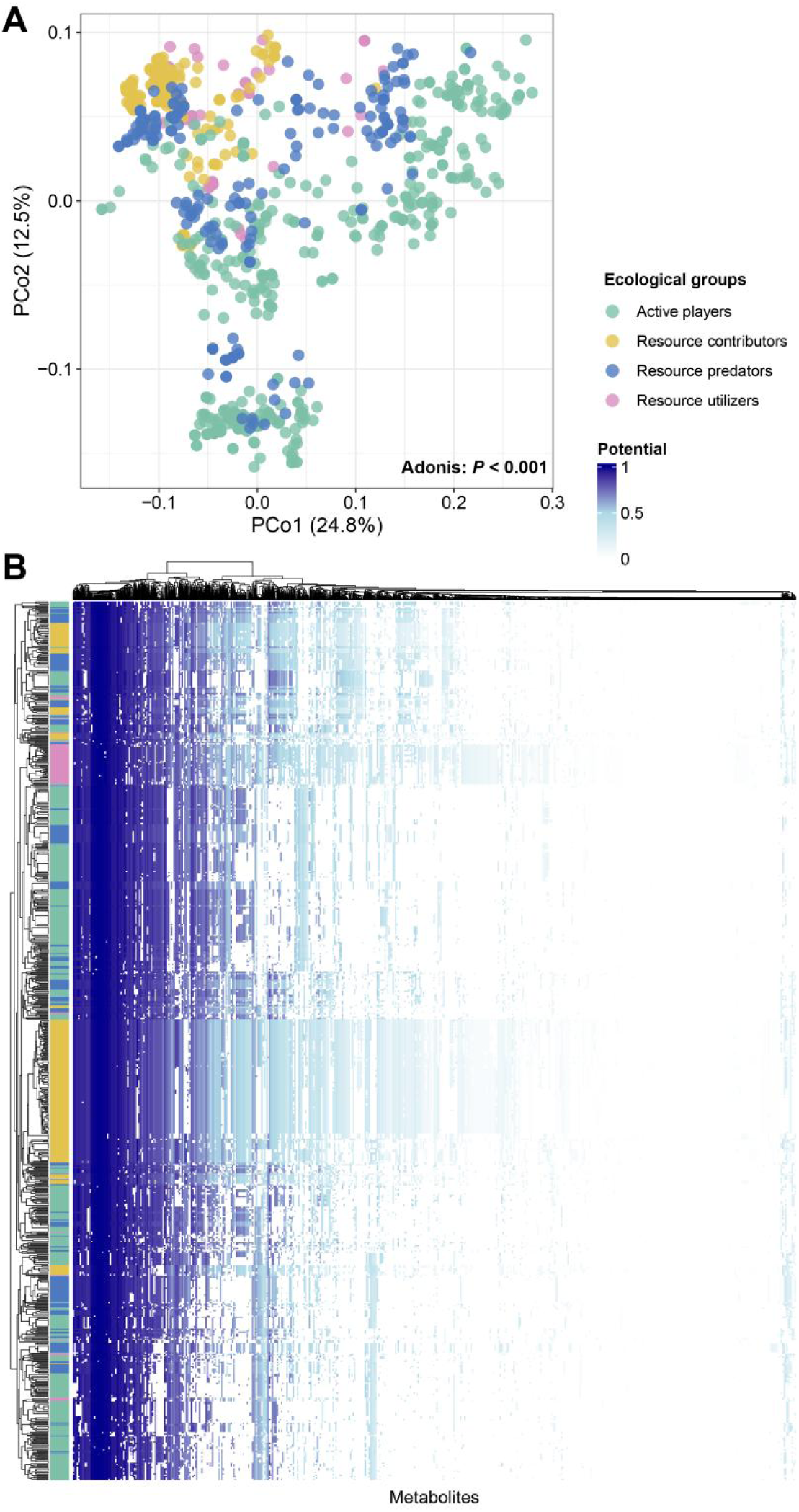
Distinct metabolic competition profiles of four ecological groups. **A** Principal Coordinate Analysis (PCoA) of metabolic competition potential profiles for ecological groups. **B** Heatmap illustrating the specific metabolic competition potential of different ecological groups. The color scale indicates the potential, defined as the proportion of the total community with which a focal strain competes for a specific metabolite.

**Supplementary Fig. 10.**
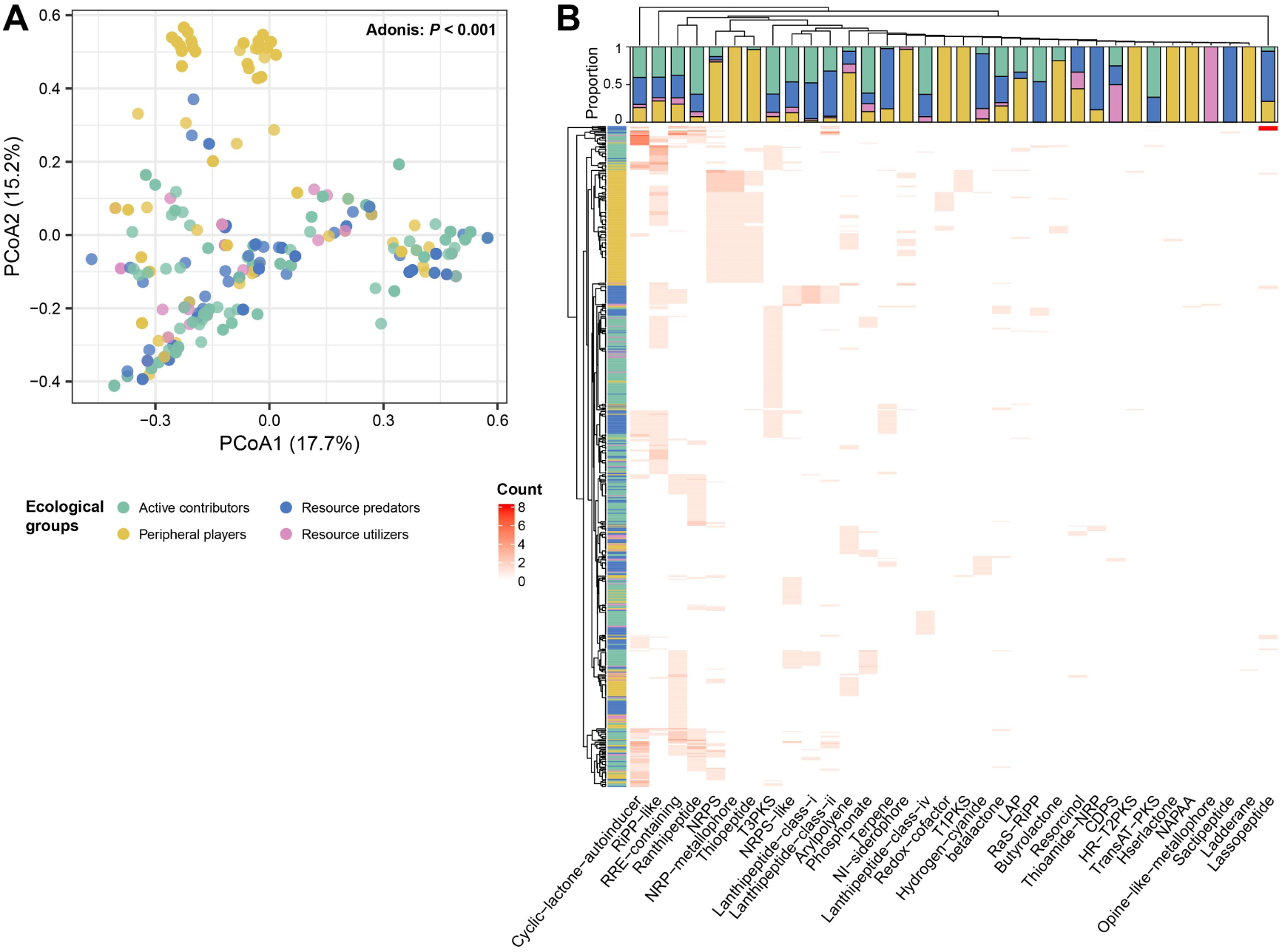
Distribution pattern of biosynthetic gene clusters (BGCs) within four ecological groups. **A** Principal Coordinate Analysis (PCoA) of BGCs for four ecological groups. **B** Heatmap showing the distribution of BGCs among genomes, and barplots illustrating the proportion of genomes containing BGCs in different ecological groups.

**Supplementary Fig. 11.**
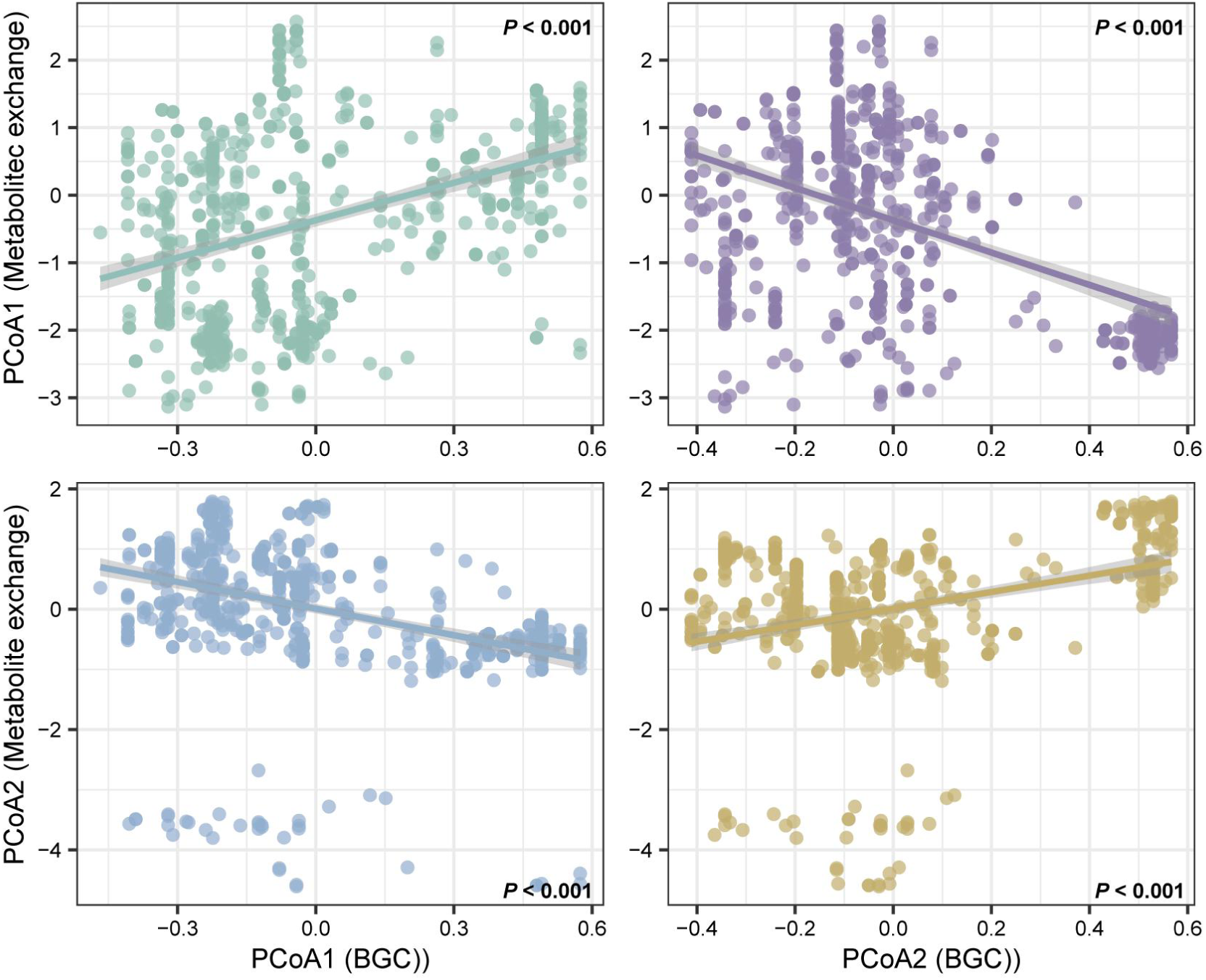
Associations of metabolite exchange with BGCs. The linear relationships of PCoA1 and PCoA2 of metabolite exchange matrix with PCoA1 and PCoA2 of the BGC matrix.

**Supplementary Fig. 12.**
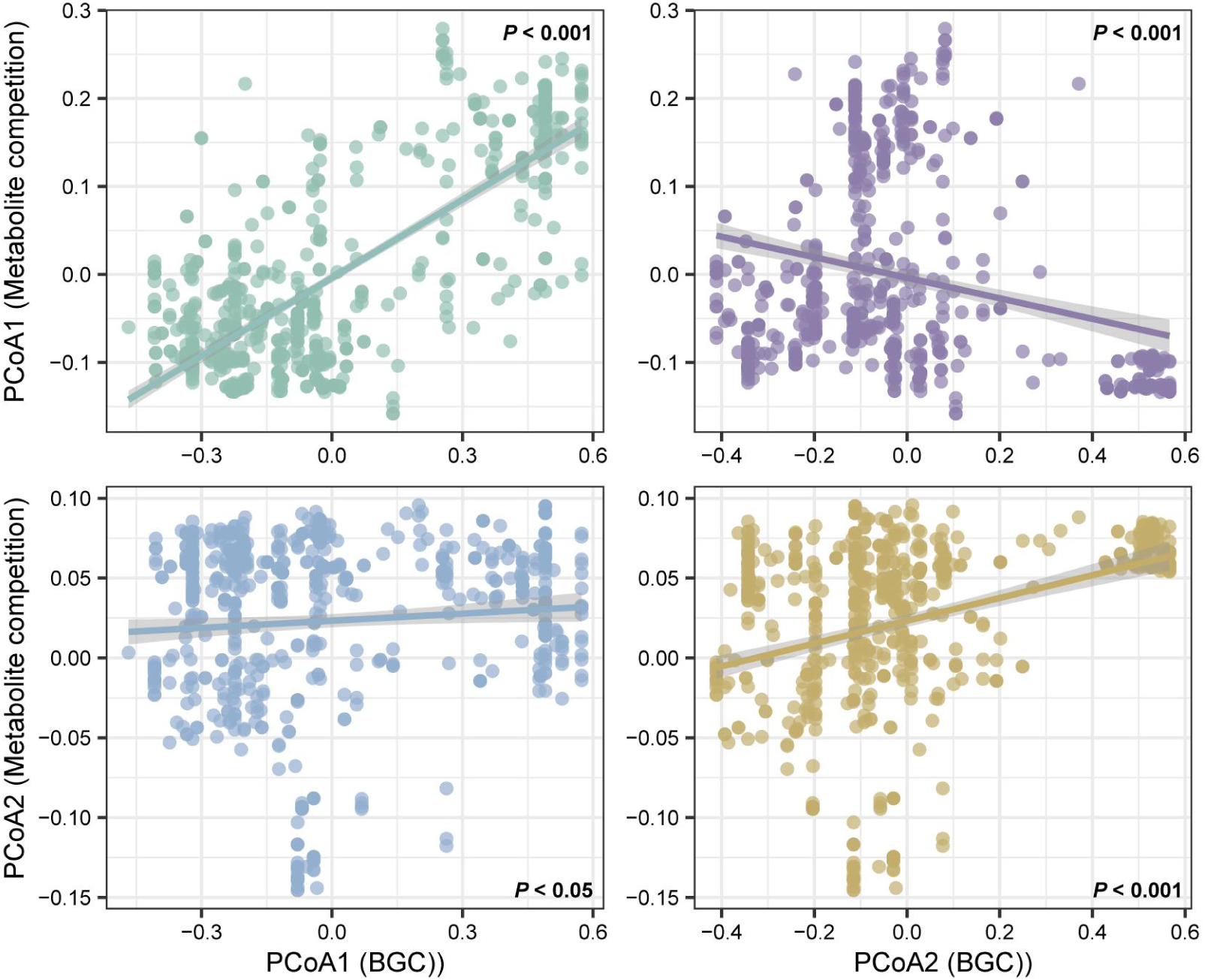
Associations of metabolite competition with BGCs. The linear relationships of PCoA1 and PCoA2 of metabolite competition matrix with PCoA1 and PCoA2 of the BGC matrix.

**Supplementary Fig. 13.**
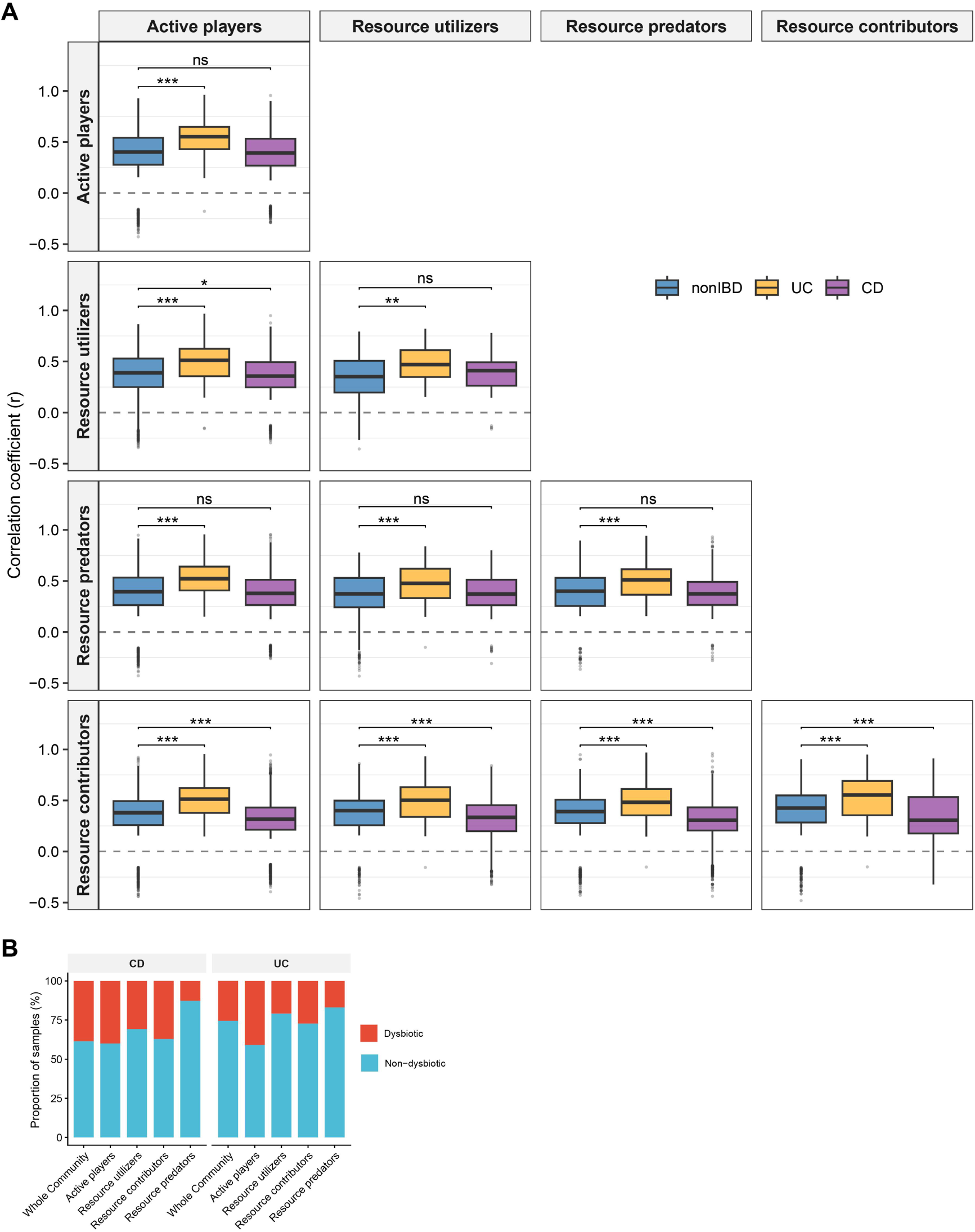
Altered ecological co-occurrence topology and group-specific dysbiosis signatures in inflammatory bowel disease (IBD). **A** Differential Spearman’s correlations within distinct ecological groups among nonIBD, ulcerative colitis (UC), and Crohn’s disease (CD). **B** Assessment of microbial dysbiosis prevalence stratified by ecological groups and the whole community. The barplots show the proportion of samples classified as dysbiotic within each ecological group. Asterisks indicate statistical significance determined by Kruskal-Wallis tests (*** *P* < 0.001, ** *P* < 0.01, and * *P* < 0.05).

**Supplementary Fig. 14.**
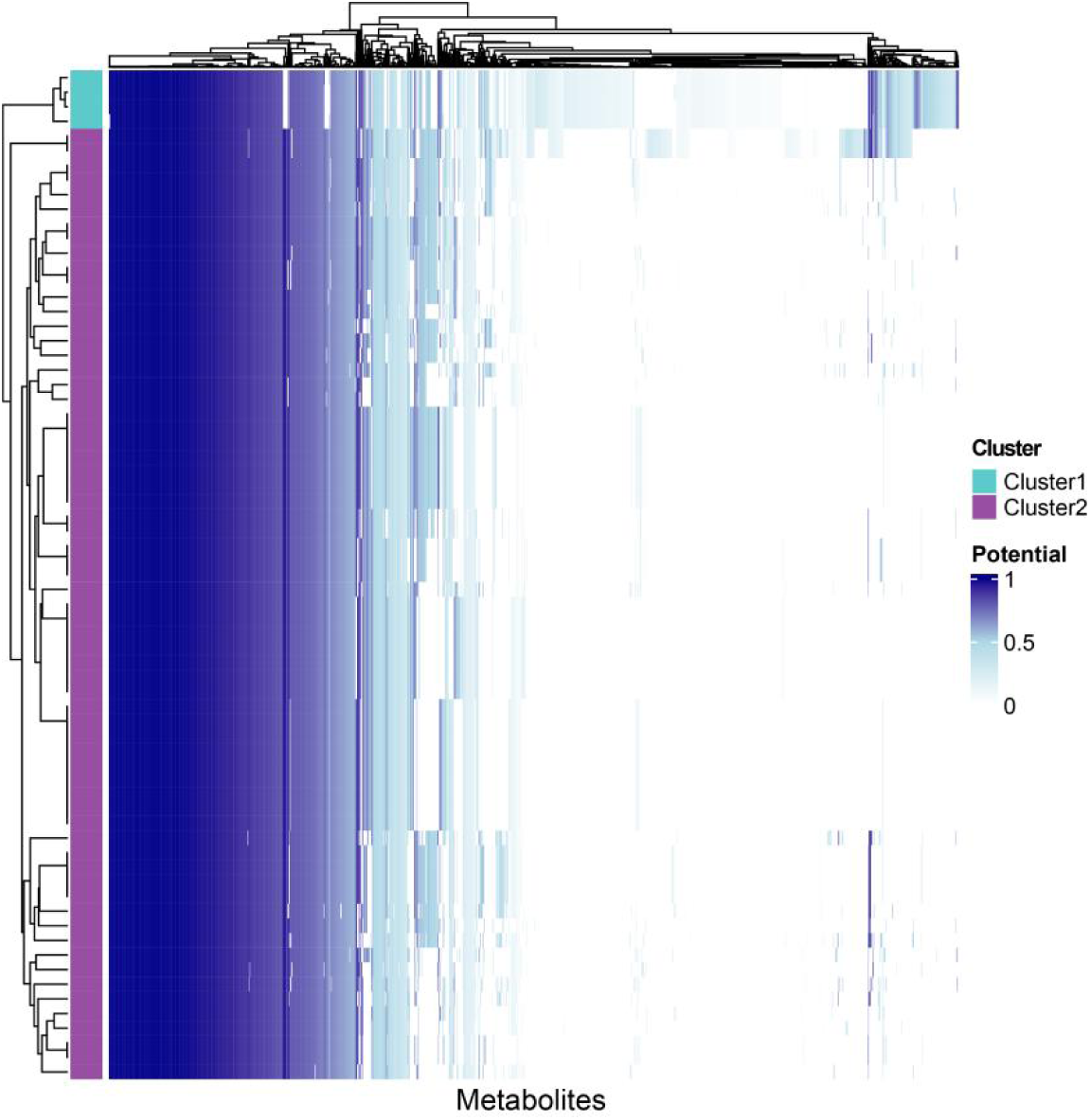
Metabolite competition between *Bifidobacterium longum* and other strains. Heatmap showing the metabolite competition between strains in cluster1 and 2 of *Bifidobacterium longum* and those from other species. The color scale indicates the potential, defined as the proportion of the total community with which a focal strain competes for a specific metabolite.

**Supplementary Fig. 15.**
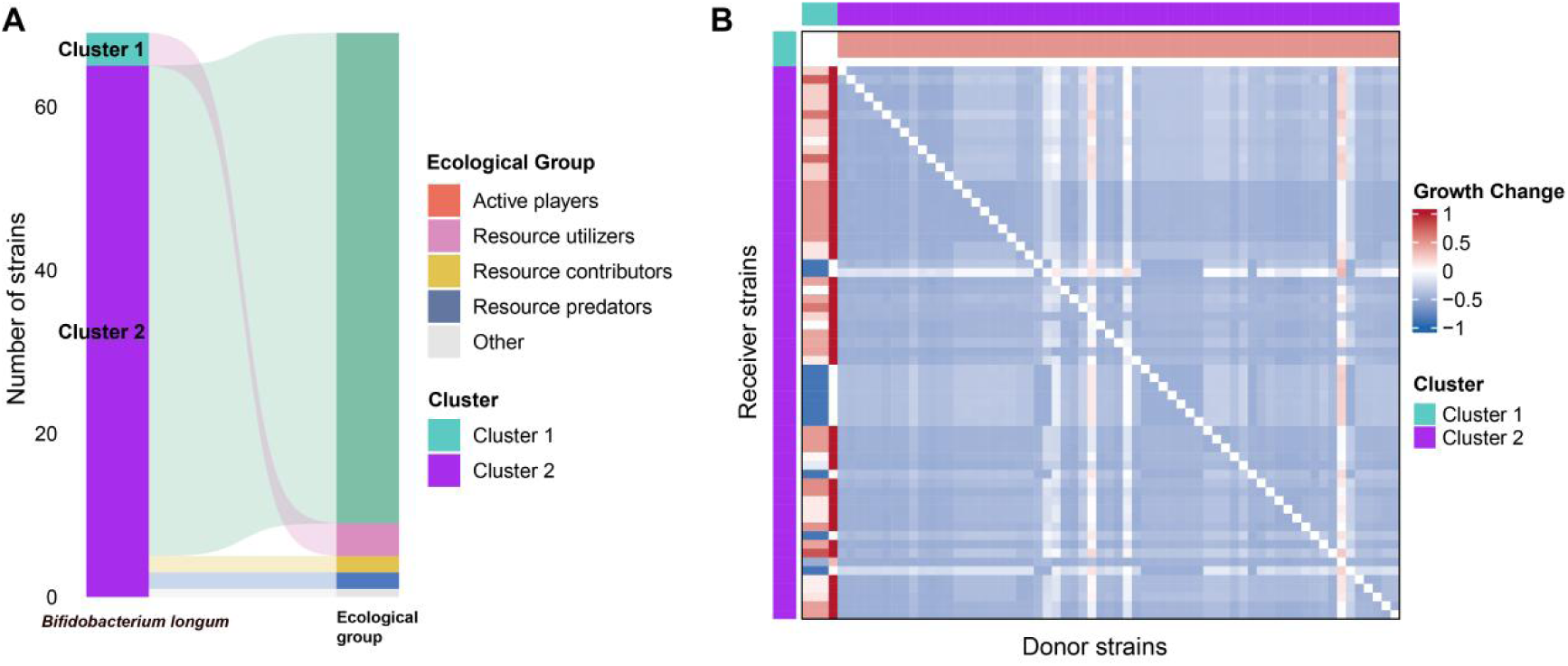
Simulated monoculture and co-culture of *Bifidobacterium longum* strains from distinct clusters. **A** The classifications of *B. longum* strains in ecological groups. **B** Heatmap displaying the relative changes in growth rates of pairwise strains under metabolic interactions compared to mono-culture conditions. The growth change was calculated as (Growth_co-culture - Growth_mono-culture) / Growth_mono-culture. Notably, rescue events (transition from zero mono-culture growth to viability) were imputed with a fixed maximum positive score (0.5) to resolve the mathematical indefinability of relative change when baseline growth is zero.

