## Supplementary material for "Complete genome-derived metabolic interactions reveal impact of gut ecology on human health": SFigures

### Supplementary figures

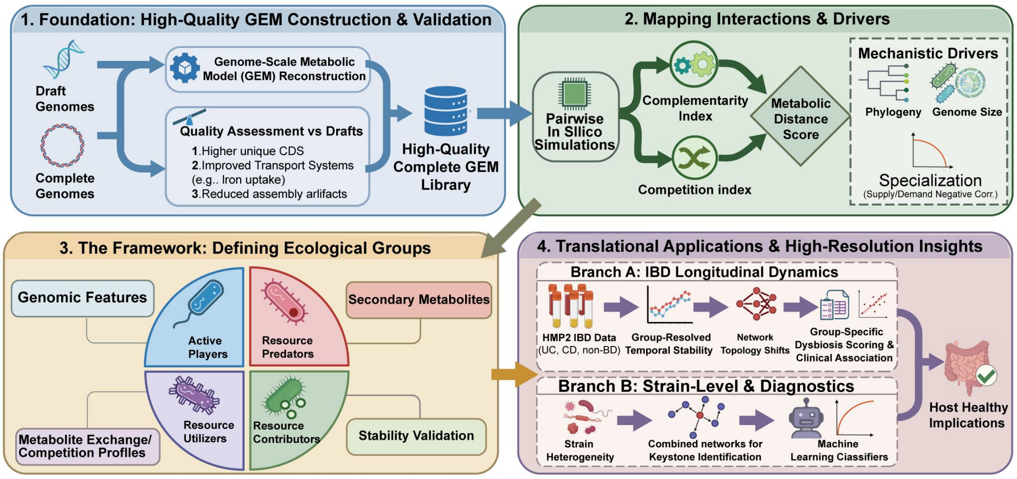


**Supplementary Fig. 1 Exploring metabolic interaction patterns and their ecological effects.** Workflow developed for metabolic interaction analysis using complete genomes.

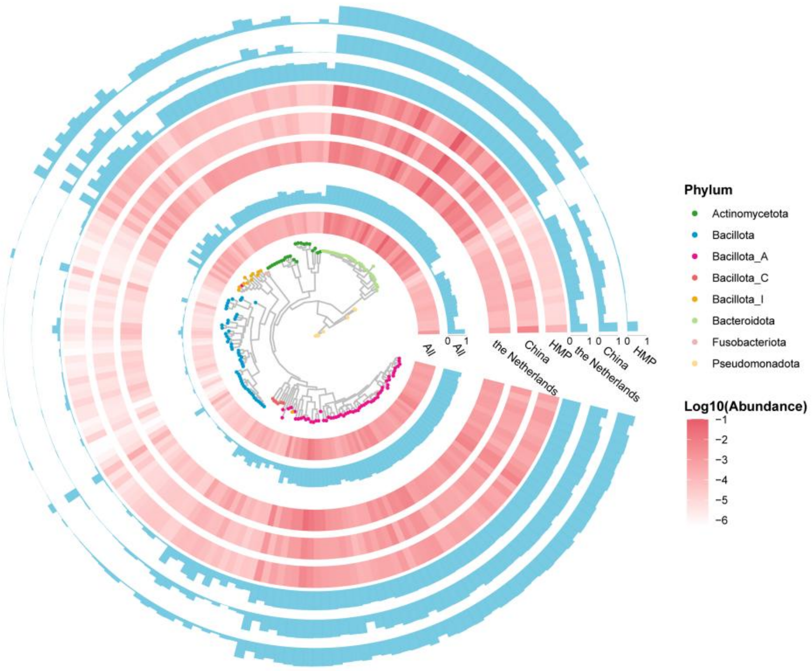


**Supplementary Fig. 2 Distributions of species in healthy individuals.** Abundance and prevalence of species in cross sectional cohorts of healthy individuals in China, HMP, and the Netherlands. The depth of the red color represents the abundance level, and the height of the blue column represents the prevalence level.

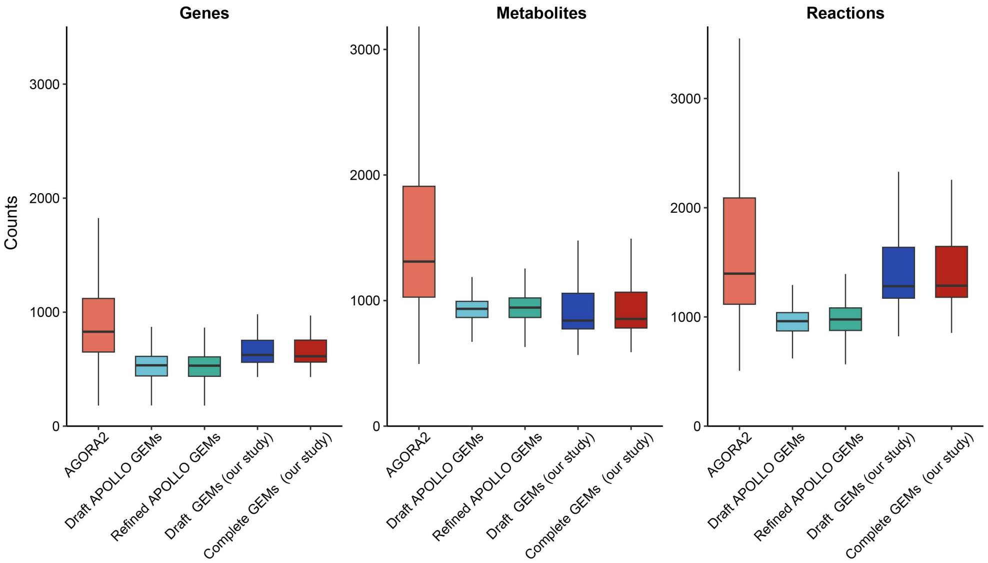


**Supplementary Fig. 3 Comparison of different GEMs databases.** The differences in the number of genes, metabolites, and reactions among different GEMs databases.

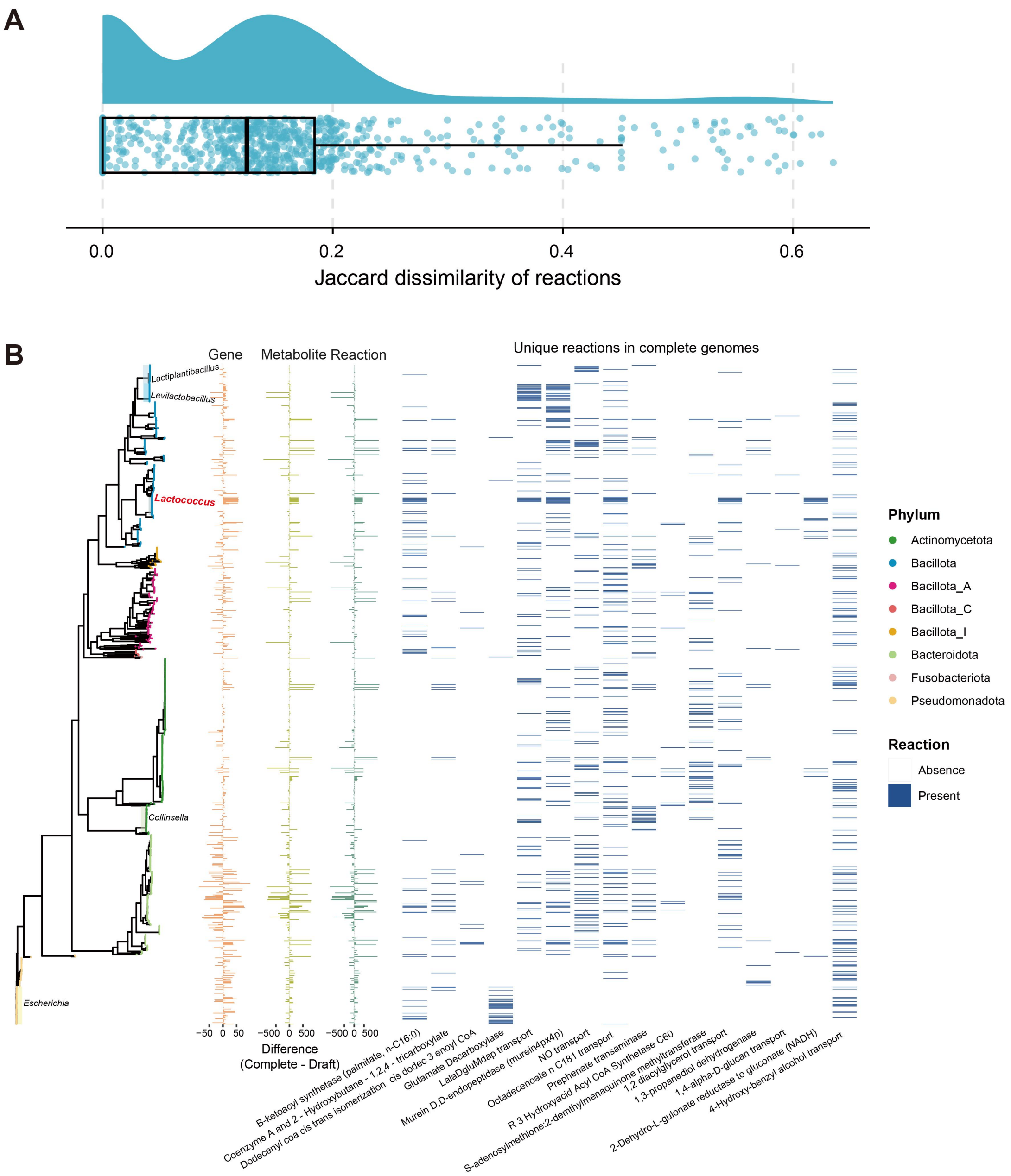


**Supplementary Fig. 4 Differences between complete and draft genome-based GEMs. A** Jaccard dissimilarity of reactions between complete and draft genome-based GEMs for each genome. **B** Variations between complete and draft genome-based GEMs. Phylogenetic tree colored by bacterial phylum is shown on the left, bar plots in the middle show the differences in the counts of genes, metabolites, and reactions for each strain, and heatmap on the right shows the unique reactions with high occurrence rates discovered in complete genome-based GEMs compared with these from draft genomes.

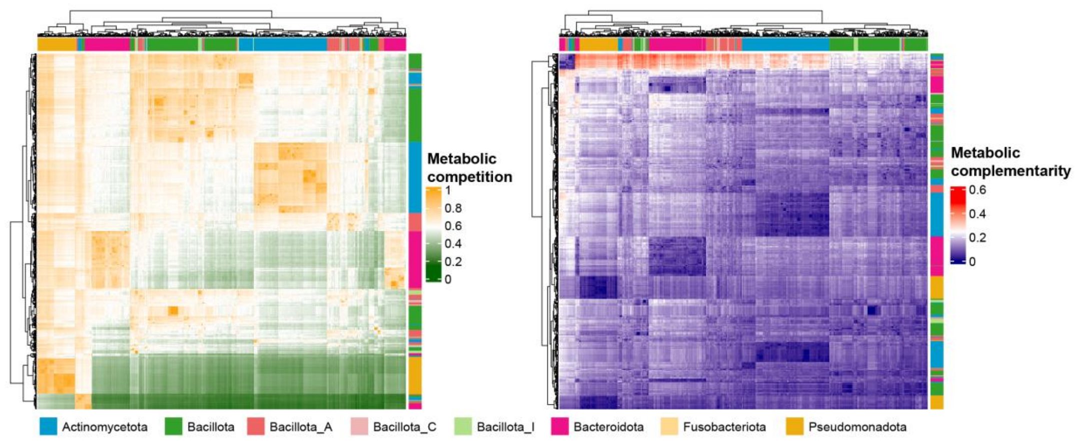


**Supplementary Fig. 5 Metabolic interaction patterns. Heatmaps showing t**he metabolic competition (**A**) and metabolic complementarity indices between every pair of genomes (**B**).

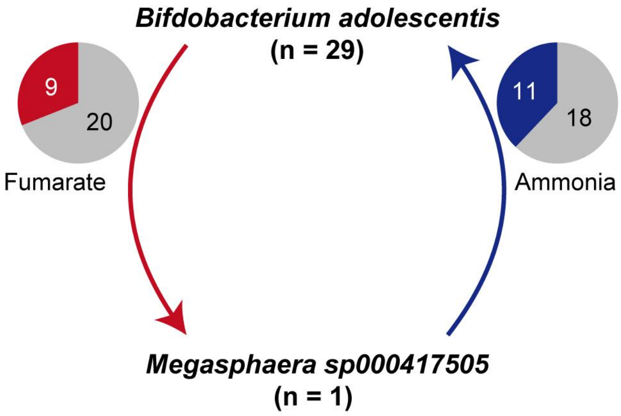


**Supplementary Fig. 6 An example of metabolite exchange.** Schematic diagram showing metabolic exchange of fumarate and ammonia between *Bifdobacterium adolescentis* and *Megasphaera sp000417505*. Pie plots show how many genes in *Bifdobacterium adolescentis* could be involved in these exchanges.

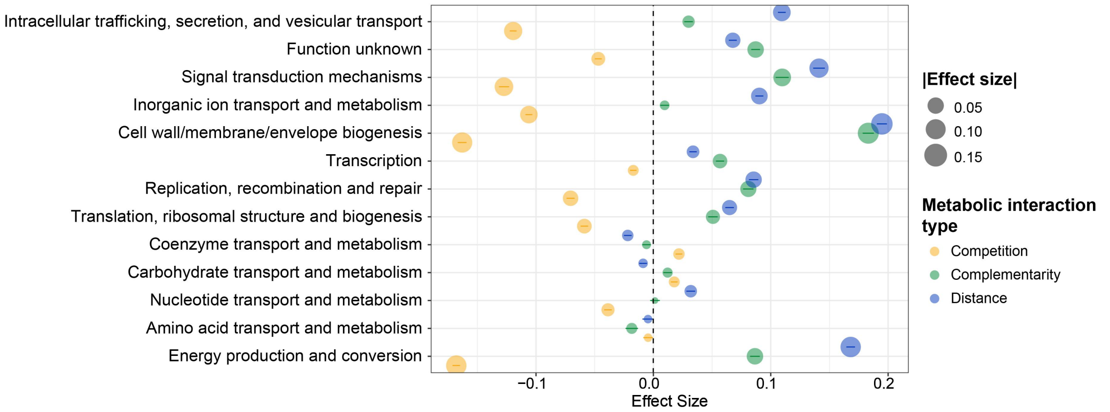


**Supplementary Fig. 7 Impacts of bacterial function on metabolic interaction.** Effect of COG biological function proportion on metabolic competition, complementarity, and distance using linear mixed-effects models. Bar length and error bar indicate the mean values and standard errors of the estimated effect sizes.

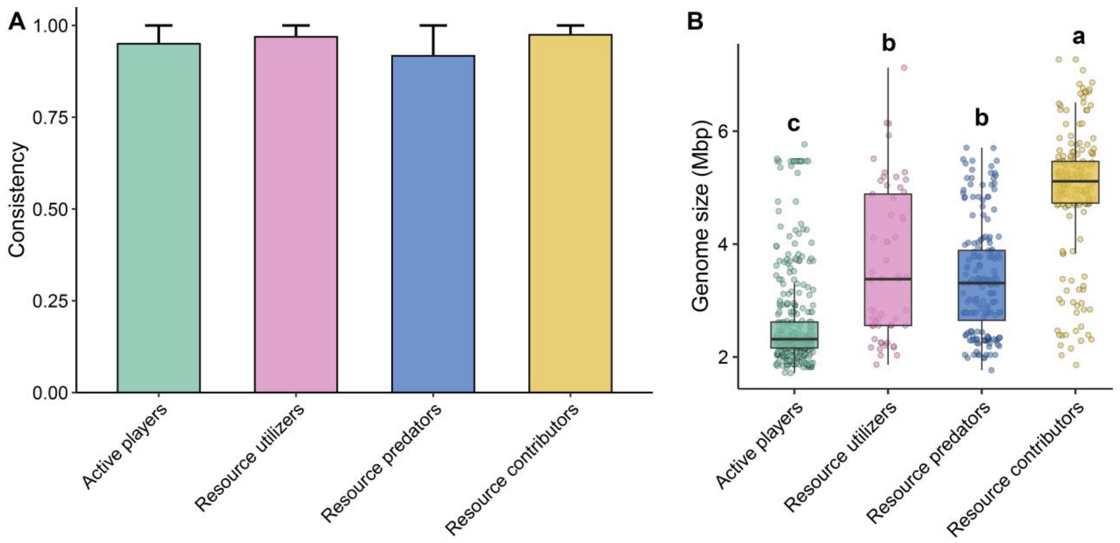


**Supplementary Fig. 8 Robustness validation of ecological group assignments and genome sizes of groups. A** To assess the stability of the inferred ecological roles (Active players, Resource utilizers, Resource predators, and Resource contributors), a bootstrap analysis was performed with 100 iterations. The skewness towards high consistency scores confirms that the classification of ecological groups is statistically robust. **A** Boxplots showing the genome sizes of strains across the four groups.

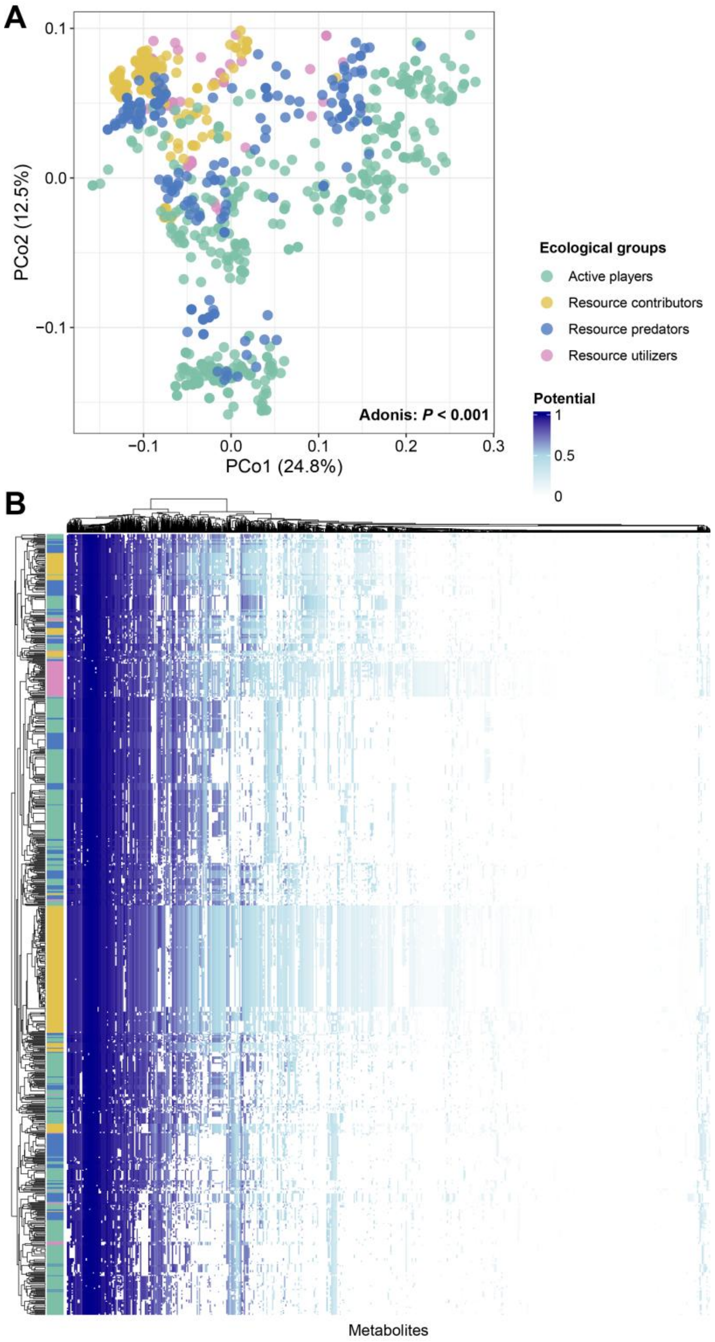


**Supplementary Fig. 9 Distinct metabolic competition profiles of four ecological groups.** **A** Principal Coordinate Analysis (PCoA) of metabolic competition potential profiles for ecological groups. **B** Heatmap illustrating the specific metabolic competition potential of different ecological groups. The color scale indicates the potential, defined as the proportion of the total community with which a focal strain competes for a specific metabolite.

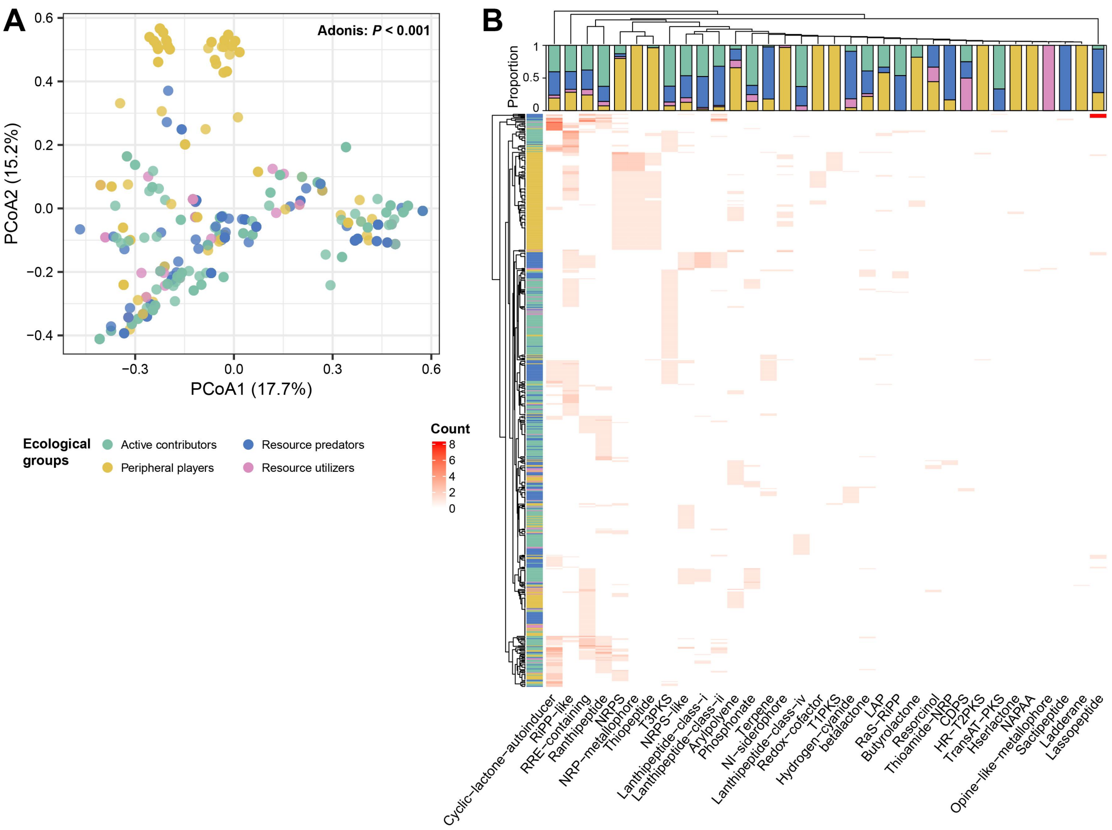


**Supplementary Fig. 10 Distribution pattern of biosynthetic gene clusters (BGCs) within four ecological groups.** **A** Principal Coordinate Analysis (PCoA) of BGCs for four ecological groups. **B** Heatmap showing the distribution of BGCs among genomes, and barplots illustrating the proportion of genomes containing BGCs in different ecological groups.

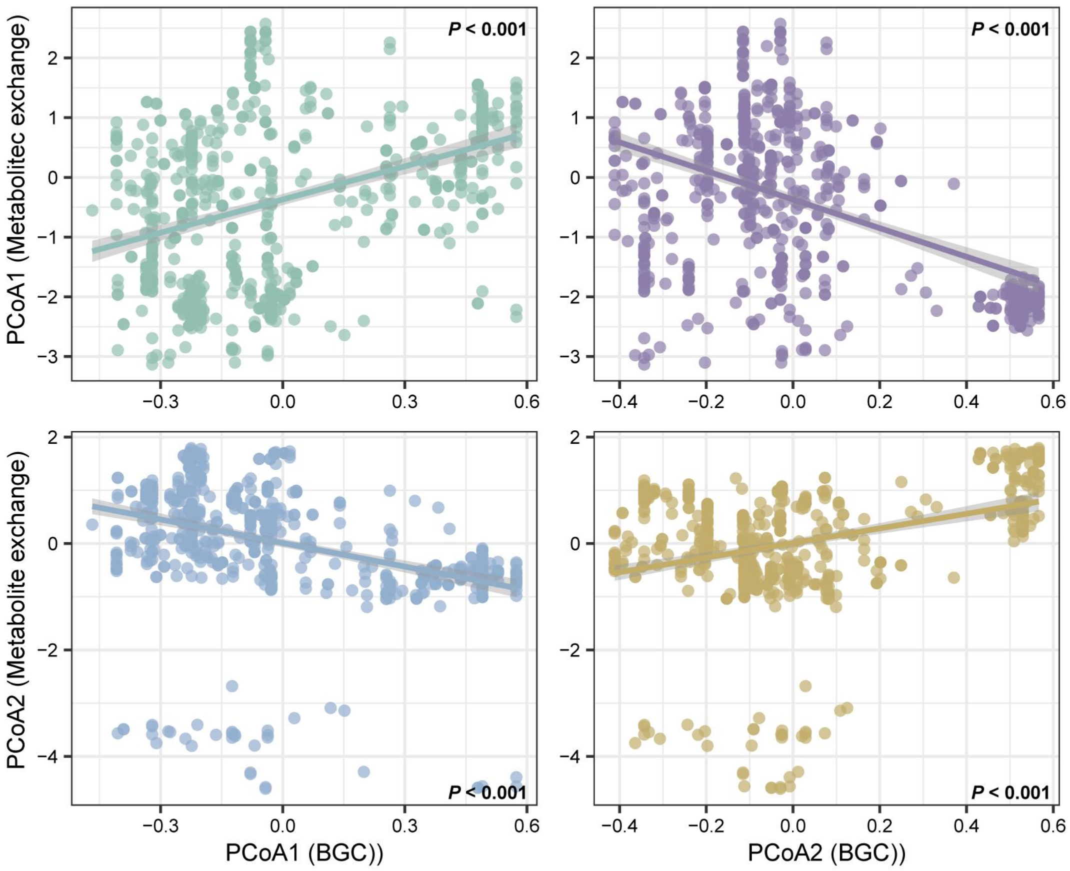


**Supplementary Fig. 11 Associations of metabolite exchange with BGCs.** The linear relationships of PCoA1 and PCoA2 of metabolite exchange matrix with PCoA1 and PCoA2 of the BGC matrix.

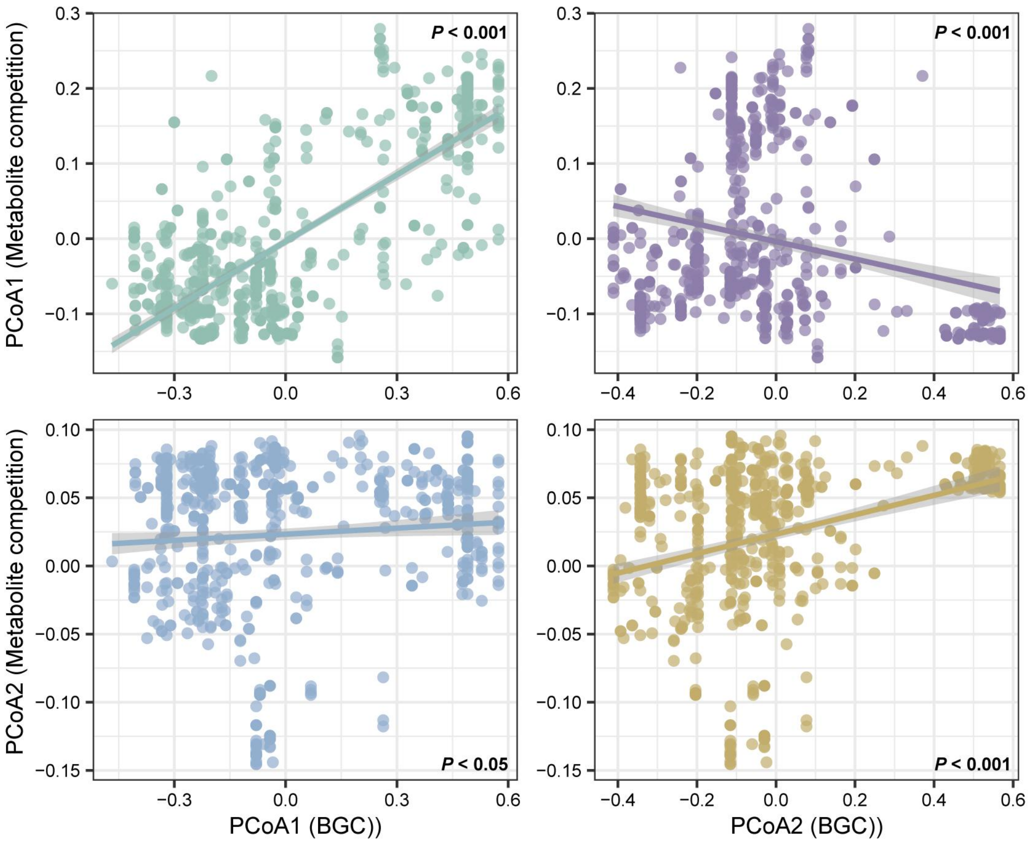


**Supplementary Fig. 12 Associations of metabolite competition with BGCs.** The linear relationships of PCoA1 and PCoA2 of metabolite competition matrix with PCoA1 and PCoA2 of the BGC matrix.

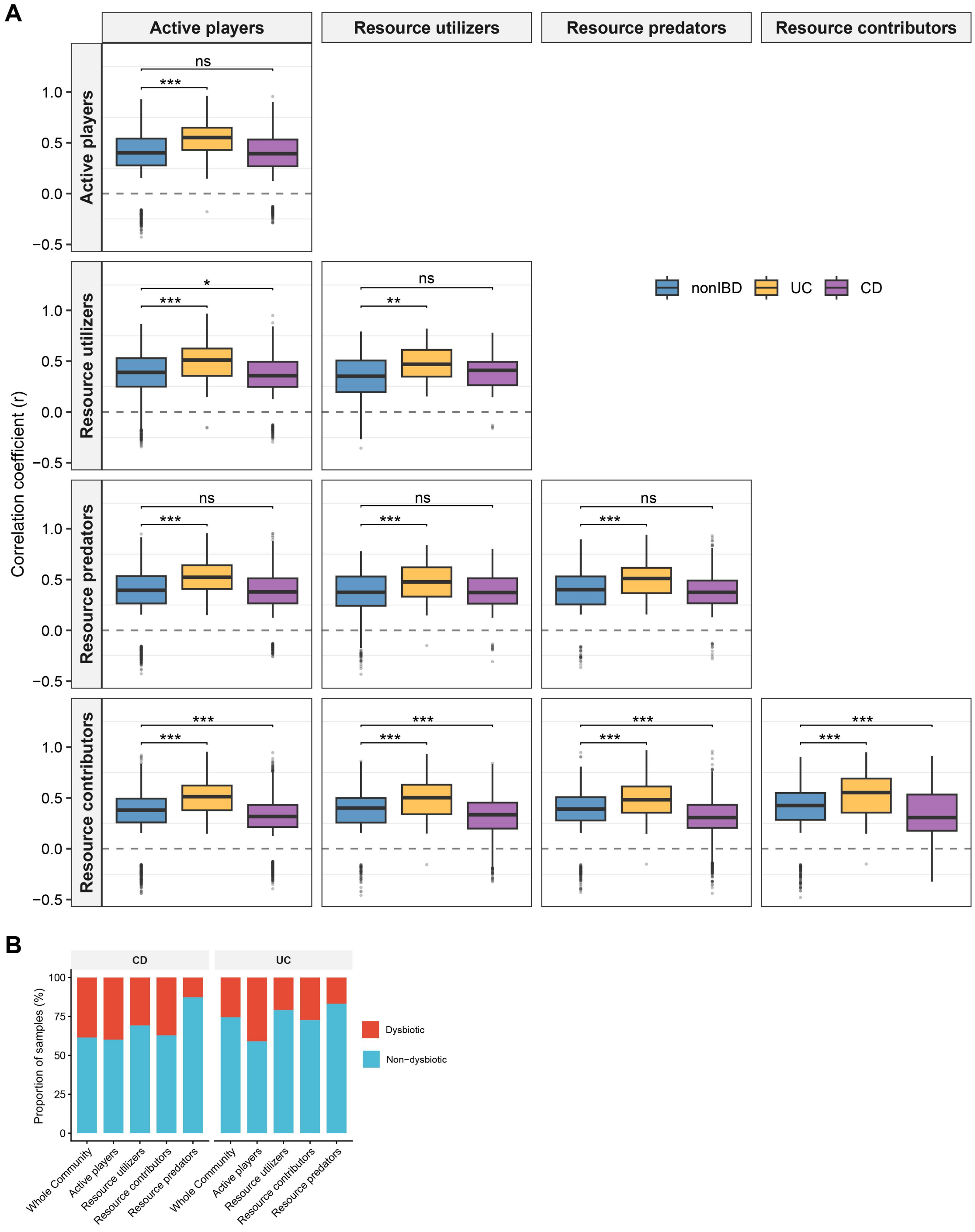


**Supplementary Fig. 13 Altered ecological co-occurrence topology and group-specific dysbiosis signatures in inflammatory bowel disease (IBD). A** Differential Spearman's correlations within distinct ecological groups among nonIBD, ulcerative colitis (UC), and Crohn's disease (CD). **B** Assessment of microbial dysbiosis prevalence stratified by ecological groups and the whole community. The barplots show the proportion of samples classified as dysbiotic within each ecological group. Asterisks indicate statistical significance determined by Kruskal-Wallis tests (*** $P < 0.001$, ** $P < 0.01$, and * $P < 0.05$).

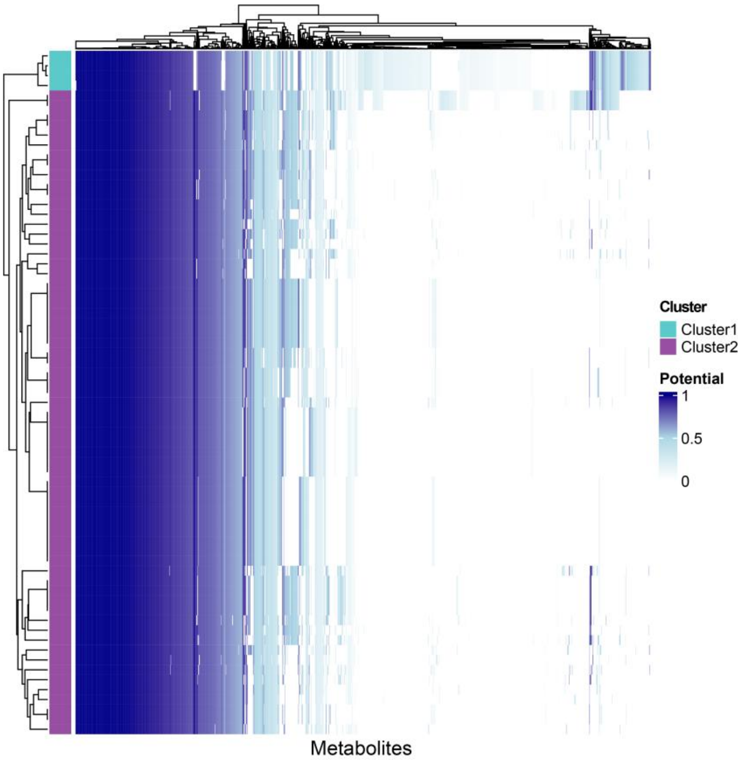


**Supplementary Fig. 14 Metabolite competition between *Bifidobacterium longum* and other strains.** Heatmap showing the metabolite competition between strains in cluster1 and 2 of *Bifidobacterium longum* and those from other species. The color scale indicates the potential, defined as the proportion of the total community with which a focal strain competes for a specific metabolite.

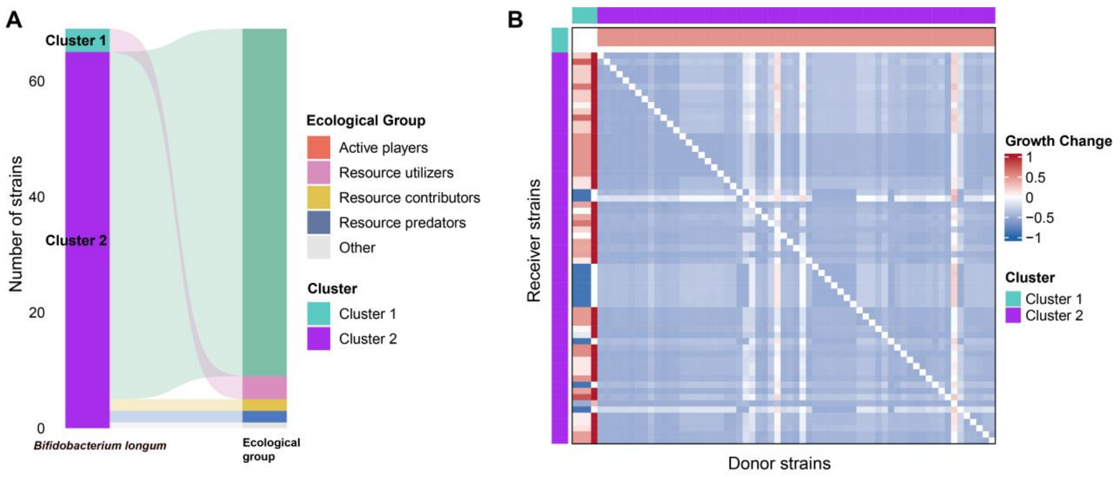


**Supplementary Fig. 15 Simulated monoculture and co-culture of *Bifidobacterium longum* strains from distinct clusters. A** The classifications of *B. longum* strains in ecological groups. **B** Heatmap displaying the relative changes in growth rates of pairwise strains under metabolic interactions compared to mono-culture conditions. The growth change was calculated as (Growth_co-culture - Growth_mono-culture) / Growth_mono-culture. Notably, rescue events (transition from zero mono-culture growth to viability) were imputed with a fixed maximum positive score (0.5) to resolve the mathematical indefinability of relative change when baseline growth is zero.
